# Prevalence of sympathetic fibers within the rat cervical vagus, and functional consequence on physiological effects mediated by vagus nerve stimulation (VNS)

**DOI:** 10.1101/2025.05.21.655349

**Authors:** Ashlesha Deshmukh, Rex Chin-Hao Chen, Justin Chin, Bruce Knudsen, James Trevathan, Andrew J. Shoffstall, Kip Ludwig

## Abstract

**Introduction:** Electrical stimulation of the vagus nerve (VNS) is an FDA approved therapy for epilepsy, depression and rehabilitation after stroke, with recent clinical trials to treat heart failure and inflammation. VNS is often assumed to activate either parasympathetic efferents projecting to visceral organs, and/or sensory afferents projecting from these organs, for its therapeutic effects. Recent studies in humans, swine and dogs have shown that sympathetic nerve fibers from the sympathetic trunk (ST) can frequently be found within the cervical vagus nerve (VN). However, the prevalence and functional consequence of sympathetic fibers on VNS have yet to be elucidated in the most common high throughput animal model to study disease, the rodent.

**Methods:** We carefully traced ST from sympathetic cervical ganglion (SCG) to find its location in the carotid sheath with reference to the VN in a cohort of Long Evans rats. We then assessed the prevalence of ST fibers with the cervical VN across the cohort using microCT and immunohistochemistry. Finally, we stimulated the VN and the ST in isolation, and where they were conjoined, to evaluate the ST contribution to changes in heart rate. VNS induced heart rate changes are a commonly used surrogate for changes in sympathetic/parasympathetic tone.

**Results:** The ST frequently runs in very close proximity to the VN in rats when traced caudally from the SCG. The ST is even conjoined with the VN for stretches within the carotid sheathe at the most common location to place an epineural cuff. Cross-connecting branches were found between the ST and the VN.

VNS performed at locations where there was minimal ST crossover induced dose-dependent bradycardia (decrease in heart rate) across the cohort, with detectable bradycardia across the cohort beginning at 50 μA (n=8 right, n=3 left). Conversely, stimulation of the isolated ST induced tachycardia (increase in heart rate) across the cohort beginning at ∼200 μA (n=7 right, n=3 left).

**Conclusion:** These data suggest that studies of VNS in the rodent model may also be stimulating sympathetic fibers from the ST in addition to canonical VN pathways. Concurrent sympathetic activation has profound implications for dissecting mechanisms of VNS for a host of diseases/disorders. As such, careful post-mortem assessment of the presence of ‘hitchhiking’ sympathetic fibers within the VN is critical for understanding sources of variability in VNS outcomes.

## Introduction

Electrical stimulation of the cervical vagus nerve (VN) - otherwise known as vagus nerve stimulation (VNS) - has been approved by the Food and Drug Administration (FDA) for human use since 1997, and implanted in well over 125,000 patients spanning 50 countries (Grasl et al., 2021; Purser et al., 2018). The ease of implantation and diversity of function of the VN make it an attractive target to address a wide variety of diseases and disorders. Originally approved for the treatment of drug-resistant epilepsy, VNS has more recently been approved for treatment of depression, obesity, and stroke rehabilitation (Food and Drug Administration, n.d.-a, n.d.-b; Handforth et al., 1998; Morris et al., 2013; Wheless et al., 2018)] There are also ongoing VNS clinical trials to treat a number of diseases/disorders, including rheumatoid arthritis, bronchoconstriction, and heart failure (De Ferrari et al., 2017; Hoffmann et al., 2012; Koopman et al., 2016; Tyler et al., 2017).

The VN consists of over 100,000 nerve fibers, including parasympathetic efferent fibers that innervate the majority of visceral organs, as well as the sensory afferent fibers from these same organs. These sensory afferents project to the nucleus of the solitary tract (NTS) in the brainstem (Dorr & Debonnel, 2006; Groves & Brown, 2005; Hoffman & Kuntz, 1957; Hoffman & Schnitzlein, 1961). NTS has connectivity to the locus coeruleus, nucleus basalis, the ventral tegmental area, and dorsal raphe nucleus, which influence noradrenergic, cholinergic, dopaminergic, and serotonergic input to the rest of the brain, respectively (Bailey et al., 2008; Davis et al., 2004; Evans & Murray, 1954; Shipley, 1982; Van Der Kooy et al., 1984; Van Giersbergen et al., 1992). In addition, NTS also projects to nucleus ambiguous and the caudal ventral lateral medulla, which increases parasympathetic (rest and digest) and decreases sympathetic (fight or flight) outflow to the body (Davis et al., 2004; Kitchen et al., 2006; Scislo & O’Leary, 1998).

Not surprisingly given its diversity of function, there is a growing body of evidence that VNS dose response curves for a specific therapeutic outcome are quite complex. For example, large animal studies investigating the effects of VNS on heart rate (HR) intended to treat cardiac dysfunction demonstrate that lower levels of stimulation induce tachycardia. However, at increasing levels of stimulation this transitions to no HR effect and then bradycardia (Ardell et al., 2017; Yamakawa et al., 2015; Yoo et al., 2013). Similarly, in rodent studies to facilitate brain plasticity to aid rehabilitation after injury, an ‘inverted U curve’ in therapeutic effect is often observed. Specifically, the effectiveness of VNS in promoting rehabilitation plateaus at (400-800 µA) and actually decreases at higher amplitudes (Borland et al., 2016; Buell et al., 2018; Manta et al., 2009). More recently, in rodent studies of VNS to treat inflammation, VNS has been shown to increase and decrease pro-inflammatory molecules depending on current dose (Tsaava et al., 2020).

One possible contributor to these complex dose response curves of VNS is the recent finding in canines, pigs, and humans that the cervical VN frequently contains sympathetic fibers in addition to parasympathetic fibers (Jayaprakash et al., 2023; Kronsteiner et al., 2024a; Onkka et al., 2013; Ruigrok et al., 2023; Seki et al., 2014; Verlinden et al., 2016; Yuan & Silberstein, 2016). Depending on model, these sympathetic fibers can occur due to cross connections between ST and VN, or due to the ST itself running alongside and conjoined with the cervical VN within the carotid sheath (De Burgh Daly & Hebb, 1952; Franco-Riveros et al., 2024; Kronsteiner et al., 2024a; Seki et al., 2014; Settell et al., 2020).

Despite recent evidence of sympathetic fibers within the VN across multiple large animal models, their presence and functional implications on VNS have yet to be assessed in the rodent model. The rodent model is the most common animal model for VNS to treat diseases/disorders, as it allows for more cost-effective and higher throughput studies to achieve appropriate statistical power. Previous rodent studies have reported that stimulating the ST in insolation causes tachycardia (Kawada et al., 2019), putatively through descending efferent pathways innervating the heart. In this study, we first use a combination of microanatomical dissection, histology and micro-computer tomography (microCT) to characterize the prevalence and anatomical sources of sympathetic fibers within the rodent cervical vagus. We then assess the functional implications of these sympathetic fibers on VNS induced changes in heart rate; changes in heart rate are commonly used surrogate for changes in sympathetic/parasympathetic tone. Finally, we outline the best practices for identifying and mitigating possible sympathetic activation that should be considered in the interpretation of acute and chronic VNS rodent studies.

## Materials and Methods

### Animals

All experiments were carried out according to the Institutional Animal Care and Use Committee (IACUC) approved protocols. All animals were housed according to the UW Madison Research Animal Resources and Compliance (RARC) protocol. A total of 20 adult Long Evans cadavers (n=10 right vagus nerve cutdown, n=10 left vagus nerve cutdown) were used in microdissection studies. These numbers were chosen to account for the variability in anatomy between individual animals. Of these 20 animals, a subset were first used for functional studies (average weight: 455 ± 94 g, n = 8 right, n = 3 left) to understand the effects of VNS and sympathetic nerve trunk stimulation (STS) on HR.

### Anesthesia

Isoflurane (2%) mixed with oxygen (1 L/h flow rate) was used an inhalant to induce anesthesia for animal sedation. Buprenorphine (Concentration: 0.3 mg/mL, Dosage: 0.05 mg/kg) was delivered subcutaneously near the incision site before the start of surgery as an analgesic. Isoflurane was maintained at 2% concentration with oxygen during surgical cutdown. It was then lowered to 1 - 1.2% during the collection of functional data (Figure 6C), minimizing effects of anesthesia on autonomic responses(Kato et al., 1992; Verardo et al., 2025), while maintaining sufficient anesthetic depth.

Anesthetic depth was verified every 15 minutes, by the absence of a pain response to a toe pinch. HR was monitored throughout the experiment. A heating pad set to 37 °C was placed beneath the animals to maintain core body temperature.

### Surgical technique

#### Carotid artery and VN exposure

The subjects were placed in a supine position and the carotid artery was exposed via a midline incision, as previously documented (Chang et al., 2020; Deshmukh et al., 2020; Lim et al., 2016; Noller et al., 2019; Sabetian et al., 2021). Briefly, the midline incision was extended from the sternum to the mandibular angle [Figure 1A]. Then, the submaxillary gland lobes were separated along with connective tissue to expose the muscles underneath [Figure 1B]. Next, the Sternohyoid (SH) and sternomastoid (SM) muscles were blunt dissected (the **right** SM and SH were dissected to expose the **right** carotid and VN bundle while the **left** SM and SH were dissected to expose the **left** carotid and VN bundle) using cotton swabs [Figure 1C]. This exposed the carotid artery bundle with omohyoid muscle (OH) running across the surgical window [Figure 1C]. The OH was then retracted or, in some cases, cut to expose the carotid bundle [Figure 1D]. The carotid bifurcation was identified by following the common carotid artery (CCA) cranially, starting from the OH to the mandibular angle [Figure 1F]. Care was taken to avoid severing nerve connections to the parotid gland, lymph nodes and adjacent muscles [Figure 1E].

**Figure 1:**
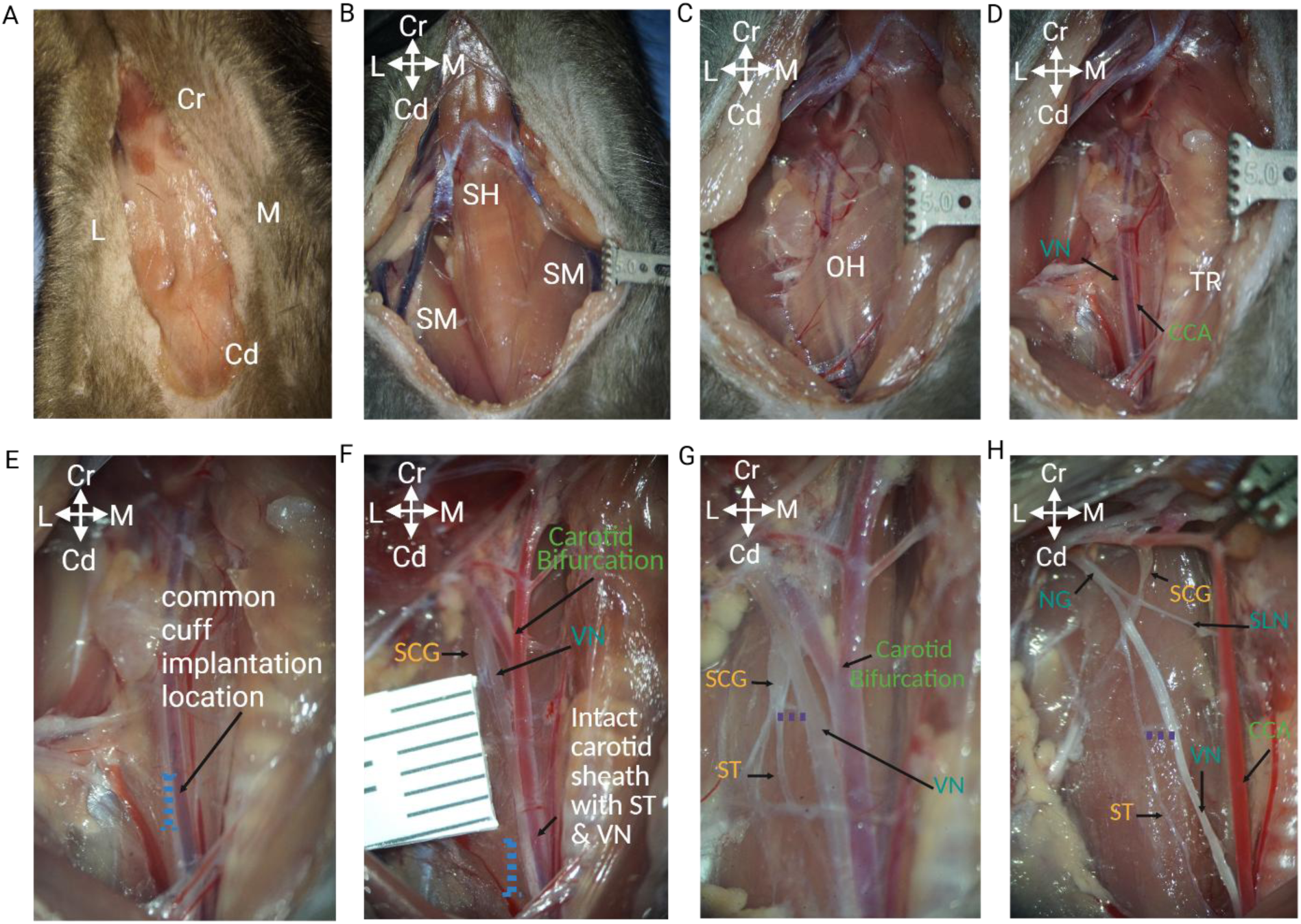
Procedure to isolate Superior cervical ganglia (SCG), sympathetic trunk (ST) and cervical vagus nerve (VN). **A)** Ventral region of the neck with the surgical pocket extending from sternum to bend in the jaw with exposed mandibular glands. Cr - Cranial, Cd – Caudal, L - Lateral, M – Medial. Convention is the same in all remaining panels. **B)** Retracted mandibular glands and exposed SH and SM muscles. **C)** Retraction along the boundary between SH and SM with exposed OH muscle running across the CCA. **D)** Retracted OH muscle with CCA and VN (nerve bundle) exposed. **E)** Common VNS cuff implantation location (blue dotted line) according to standard protocols (cranial/caudal or dorsal to retracted OH). **F)** Dissected SCG at the carotid bifurcation with ST ‘hitchhiking’ with the VN. The carotid sheath and the connective tissue are intact ∼2-3 mm caudal to the bifurcation. Cuff implantation location for reference (blue dotted line). **G)** Dissected connecting branch between caudal pole of SCG and joining the VN (purple dotted line). This branch is separate from the ST which runs parallel to the VN along the CCA. **H)** VN originating from NG. The SLN originates at the caudal pole of NG. The ST originates from the caudal pole end of the SCG. A second connecting branch was observed between ST and VN ∼5-6 mm caudal of the carotid bifurcation (purple dotted line). Sternomastoid (SM), Sternohyoid (SH), omohyoid (OH), carotid artery (CCA), superior cervical ganglia (SCG), nodose ganglia (NG), Trachea (TR), superior laryngeal nerve (SLN), vagus nerve (VN), sympathetic trunk (ST). Note: This pictorial representation is from the rodent right side. The left side cutdown pictures are reported in Supplementary figure 5 and 7. This figure was Created in https://BioRender.com.

#### Dissecting Superior cervical ganglia (SCG) and tracing the ST

Custom-made glass rods and hooks [Supplementary Figure 1] were used to minimize bleeding while handling the carotid bifurcation. The carotid bifurcation was carefully lifted and tilted laterally, providing access to the dorsal side of the bifurcation. The SCG was found where it is adhered to the dorsal side of internal carotid artery (ICA), with branches to ICA, external carotid artery (ECA), carotid body, nodose ganglia (NG) and other muscles in the vicinity [Supplementary Figure 2]. Blunt dissection was performed to isolate the SCG from the artery [Figure 1F and G]. This surgical approach was different from the established techniques which dissect and clear connective tissue at the carotid bifurcation, between the ICA and ECA (Madhani et al., 2022). This was intended to avoid damaging the carotid body and physiological sensors, or their associated nerve fibers at the carotid bifurcation [Supplementary Figure 3], that provide input to the autonomic system (Paton et al., 2013; Prabhakar, 2016). The ST was identified as the branch originating from caudal end of the SCG [Figure 1G]. All other surrounding nerves were preserved throughout the surgical procedure, including the superior laryngeal nerve (SLN), originating from nodose ganglion (NG) and crossing to the midline [Figure 1H], the accessory nerve, aortic depressor nerve (ADN) and hypoglossal nerves [Supplementary figure 3, 4]. The ST was traced from the SCG to the sternum. Its course relative to the VN, especially at the common cervical cuff implantation (cranial/caudal or dorsal to retracted OH) (Madhani et al., 2022; Noller et al., 2019; Somann et al., 2018) along with any connecting branches between the two nerves, was documented [Figure 1G, H].

### Physiological functional studies

#### Electrodes

Stimulation was delivered using custom made bipolar hook electrodes. The electrodes were fabricated using Platinum Iridium (PtIr) wire (A-M Systems, Bare 0.005” diameter, Category number: 778000). The wire tips were exposed by applying heat using thermal wire strippers to remove about 1-1.5 mm of insulation. They were bent using forceps to make a hook of ∼1 mm diameter with circumference being ∼270 degrees. The viability of the electrodes was verified prior to experiments using electrochemistry impedance spectroscopy [Supplementary methods 1]

#### Stimulation

Bipolar, charge-balanced biphasic 300 µsecs (150 μs per phase) electrical pulses were delivered at 6.25 Hz to the VN and ST individually based on prior literature (Kawada et al., 2019) using Tucker-Davis Technologies (TDT) systems [Figure 6B]. This frequency is not divisible into 60 Hz evenly and therefore 60 Hz noise is attenuated when averaging data over multiple stimulation presentations.

Amplitudes for stimulating the VN and ST were first increased using larger steps until a threshold for bradycardia or tachycardia was reached, respectively. The initial threshold for bradycardia was determined by a heart rate decrease of 10 beats per minute (BPM), whereas a heart rate increase of 5 BMP was used for tachycardia. More granular dose response curves (DRCs) were then generated by delivering stimulation at amplitudes below, at, and above threshold. In a subset of subjects (n=2, right and n=2, left), clinically relevant frequencies of 25 and 40 Hz were also used. In all cases, stimulation was delivered for 30 secs followed by 2 mins or longer to allow heart rate to recover to baseline [Figure 6C]. Finally, contributions of the VN and ST to observed HR changes were verified with nerve transections caudal to the stimulation electrodes [Figure 6B].

##### Electrode nerve interfacing

To isolate and minimize off-target activation of other nerves due to electric field spillover, the surgical pocket was dried using gauze and two pieces of **ParaFilm^®^ M Lab Film** (S- 25928, ULINE) were placed under the location of nerve-electrode interface for each nerve. The sizing and location of the film was selected to minimize interaction between the two nerve interfaces and as insulation from surrounding tissues. A drop of Kwik-Sil (World Precision Instruments, KWIK-SIL-S), a silicone adhesive, was then applied on top of the nerve interface to electrically isolate and secure the electrode to the nerve. Care was taken to ensure that Kwik-Sil did not come between the electrode and the nerve. In-vivo impedance checks were performed using TDT systems to verify that the values were below 30 kΩ. To ensure both nerves did not dry out and remained physiologically viable throughout the experiment, they were occasionally sprayed with room temperature saline using a spray bottle.

Simultaneously, extra fluid and edema built-up in the surgical pocket were removed as needed during the experiment using surgical gauze. This reduced the chances of creating current shunting pathways and off-target activation.

##### Physiological signal recording

Physiological signals were acquired using a PowerLab system (PowerLab 4/30, ADInstruments, New Zealand) using a 1 k/sec sampling rate. The electrocardiograms (ECG) were recorded using a 3-electrode set-up with needle electrodes placed on the right front paw (negative), left front paw (positive) and the tail (reference) [Figure 6A]. The ECG signals were visualized in real time using LabChart (ADInstruments, New Zealand) and filtered using a digital bandpass filter (35-200 Hz). An independent visualizing channel was used to monitor the HR, calculated by detecting the QRS complex peaks using standard deviation thresholding. Finally, the calculated HR signal was passed through a median filter with a window width of 3001 samples to smooth the signal and remove any sudden fluctuations. This filtered data was saved as another independent channel and used for quantifying HR changes. The stimulation trigger was recorded using a BNC cable as digital I/O channel. HR changes were quantified as the maximum change in HR from baseline (HR value at the start of the stimulation trigger) during stimulation.

##### Histology

Histological processing of intact cervical carotid bundle cross-sections was used to verify the presence of sympathetic fibers using immunohistochemistry (IHC). These cross-sections were collected from a separate cohort of animals and not from a consistent specific location along the cranial-caudal axis. The goal was to confirm the presence of sympathetic fibers independent of slight variations in the cuff placement locations across different studies. Samples were stained using tyrosine hydroxylase (TH+), a known marker for postganglionic sympathetic fibers (Keast, 1995a) by a team at Duke University [Supplementary methods 2]. VN parasympathetic fibers, the majority of which are non-noradrenergic (tyrosine hydroxylase negative), were used to distinguish between the two nerves in the carotid bundle. The complete protocol can be found at SPARC_Duke_Grill_OT2-OD025340_VagusNerve_IHC_TH (protocols.io)

##### MicroCT Imaging

Rat heads (n = 4 total, n = 2 sourced from UW Madison were used for pilot studies to refine imaging techniques and n=2 sourced from Case Western Reserve University used in the final analysis) were carefully disarticulated from euthanized adult Long Evans rats, post euthanasia, secondary to other approved studies. The skin was removed from the samples and the mandibular glands and sternomastoid muscles were resected. Care was taken to leave the carotid sheath and surrounding tissues in the cervical region intact. To increase the contrast of soft tissue during microCT imaging, the samples were stained with phosphotungstic acid (PTA), a radiopaque stain that binds to collagen, allowing for nervous tissue to be visualized in the scans (Thompson et al., 2019; Upadhye et al., 2022).

The samples were placed in a 50 mL falcon tube with 3% v/v PTA (Sigma-Aldrich) in deionized water. The samples were then set on a nutating shaker for two weeks, replacing the solution midway. The sample was then dehydrated via increasing concentrations of ethanol by submerging the sample in 70% ethanol for 90 minutes, replacing the solution every 30 minutes. The process was repeated with 95% ethanol.

MicroCT images were obtained using a Scanco® µCT100 instrument. The samples were imaged with the following parameters: Tube voltage (Excitation) 55 kVp, current 144 µA, integration time 500 ms, using a 0.5 mm aluminum filter. The DICOM image volumes were isotropic with a resolution of 11.4 µm [Figure 4A].

3D Visualization was performed using 3D Slicer (Kikinis et al., 2014). Image volumes were imported into the software and displayed using the “Volume Rendering” module. Low intensity voxels were omitted from visualization to reveal the sample and omit any surrounding noise. The image volume was transformed to align the axes of the sample. Segmentation of desired structures was performed by hand using the “Segmentation Editor” module in conjunction with the Slicermorph extension (Rolfe et al., 2021) for additional segmentation capabilities. The identified anatomical structures of the VN and ST were compared to microCT of dissected samples collected at UW Madison and the microdissection pictures for verification.

##### Data Analysis and Statistics

Data analysis and figures for stimulation evoked heart rate changes were generated using pyeCAP, a custom software package (https://github.com/ludwig-lab/pyeCAP, version 0.0.1) in Python 3.7. Python libraries for pandas, matplotlib, seaborn, numpy, scipy and statemodels were utilized. Statistical analysis was performed using IBM’s SPSS Statistics (version 28.0.1.0 (142)).

Functional data was collected to assess if there were any reduced bradycardic effects when stimulating the VN and the ST together where conjoined as compared to the VN alone. Wilcoxon signed-rank test was used to test if there were any statistical differences in bradycardic HR effects caused by the same stimulation parameters during VNS alone as compared to stimulating VNS+STS in conjunction (N=17 paired observations). The effect size to capture the differences in HR effects between the two stimulation groups were reported using Wilcoxon signed-rank test (a non-parametric test). Based on this calculated effect size, we derived the required sample size by power analysis to show the statistical difference between the two groups using normal_sample_size_one_tail function from the statsmodels.stats.power library in Python 3.7.

The on-target engagement during VNS and STS was verified by comparing the stimulation driven HR changes pre and post nerve transections. Normality of data for pre vs post nerve transections HR delta changes was calculated using Shapiro-Wilk test. Comparison of pre vs post nerve transections delta HR changes was performed using paired one tail t-test with bootstrapping for 1000 samples with 95% confidence intervals.

## Results

The goal of this paper was to understand the prevalence and first-order functional consequences of activating sympathetic fibers during VNS in rodents. Towards this in, in the following results we first provide an anatomical documentation across the cohort on the prevalence of the ST running joint or in proximity with the cervical VN. These data are supplemented with microCT imaging to help illustrate the relevant microanatomy in 3-dimensions without an invasive cutdown. Next, histology of cross sections of intact carotid bundle are presented to show relevance and distribution of sympathetic fibers in both the ST and the VN. Lastly, we assess the functional consequences of stimulating the ST and VN in isolation and where conjoined on heart rate, a common surrogate for VNS induced changes in sympathetic/parasympathetic tone.

### 1) ST runs in close proximity to the cervical VN in the carotid sheath

In this paper, we define the sympathovagal trunk as the conjoint bundle of the VN and the ST or a length of the VN with cross connecting fibers with the ST while the carotid bundle is defined as the nerves in the carotid sheathe running with the carotid artery. To find the sympathovagal trunk, the ST was first located as it connects to the SCG. The SCG was located dorsomedial to the carotid bifurcation in all subjects (n=10 right, n=10 left), and found adhered to the internal carotid artery (ICA) [Supplementary Figure 2].

In most animals, the ST crosses over from the medial to the lateral side of the CCA when traced from the SCG to the sternum (n=9/10, right side) [Figure 2A and B]. In some animals (n=2/10 right side), the SCG extended beyond the carotid bifurcation and was joined to the VN on the lateral side of the CCA [Figure 2C]. The ST crossover point was found to be more variable and less frequent on the left side (n=4/10).

**Figure 2:**
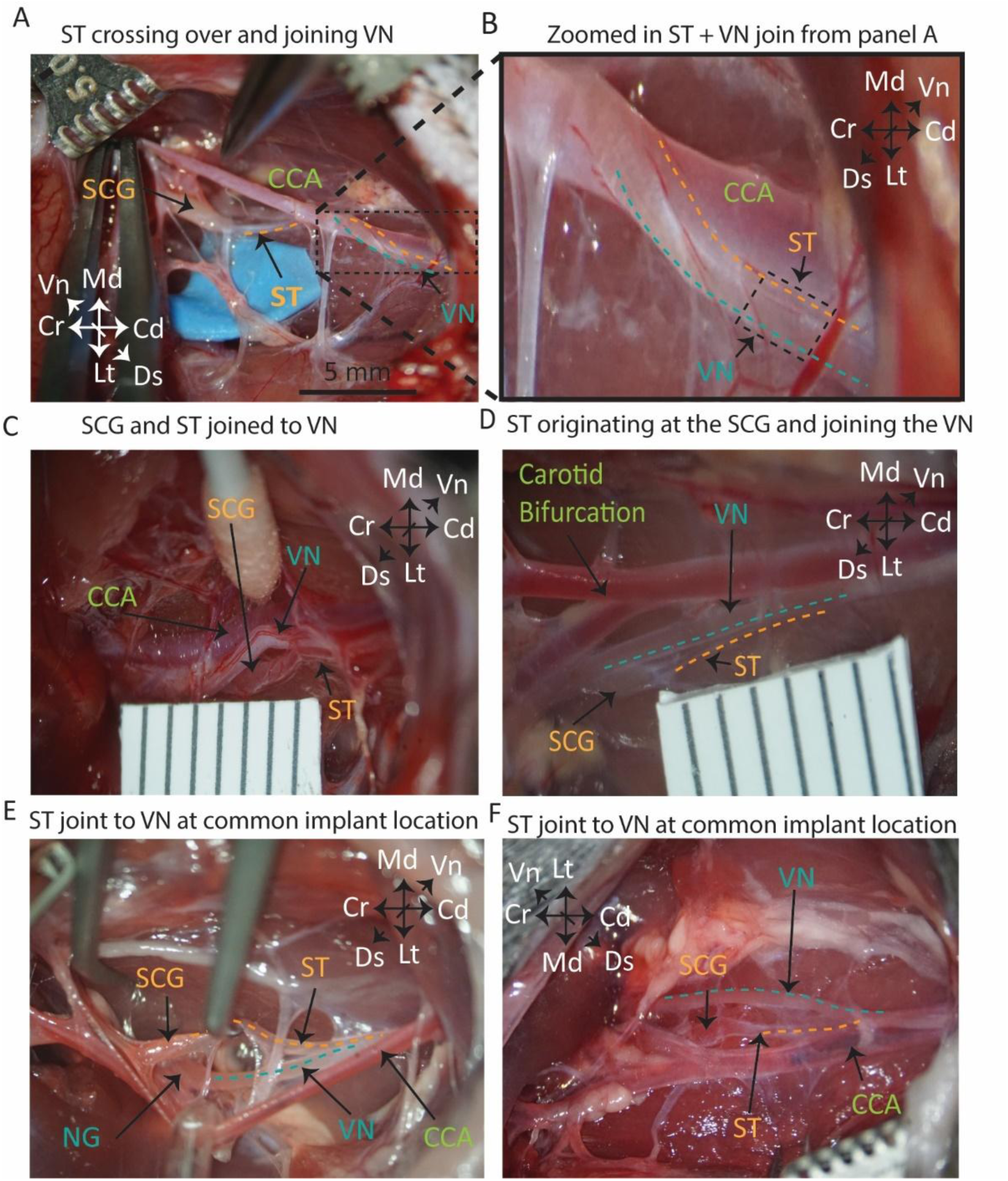
ST is joint and runs parallel to the VN at the common cuff implantation location. **A)** Anatomical visualization of ST (dark yellow dotted line) starting from caudal pole end of SCG and wrapping around the CCA from medial to lateral side to join the VN. B) Zoomed in picture of the dotted black box in Panel A; ST (dark yellow dotted line) joint and running parallel with the VN (cyan dotted line) at the common cuff implantation location. C) The SCG extending from the dorsomedial to lateral side of the CCA and joining the VN. The ST and the VN hitchhiking together and running in parallel along the CCA towards the sternum D) ST (dark yellow dotted line) and the VN (cyan dotted line) joint and running in parallel in the carotid sheath, ∼4 mm caudal to the carotid bifurcation. The undissected CCA bundle, 8- 10 mm caudal to the bifurcation, shows the ST and VN joint together. E) NG and SCG in close proximity with the carotid bifurcation retracted laterally. ST (dark yellow dotted line) originating at the SCG and the VN (cyan dotted line) originating at NG; joint as they run caudal along the CCA. F) ST (dark yellow dotted line) and VN (cyan dotted line) joint and running in parallel on the same side of the carotid artery. All microdissection pictures are from different subjects. ST: Sympathetic Trunk; SCG: Superior cervical ganglia; VN: Vagus Nerve; CCA: Common carotid artery; NG: Nodose Ganglia. Additional cohort pictures in supplementary figures 5 to 9.

**Figure 3:**
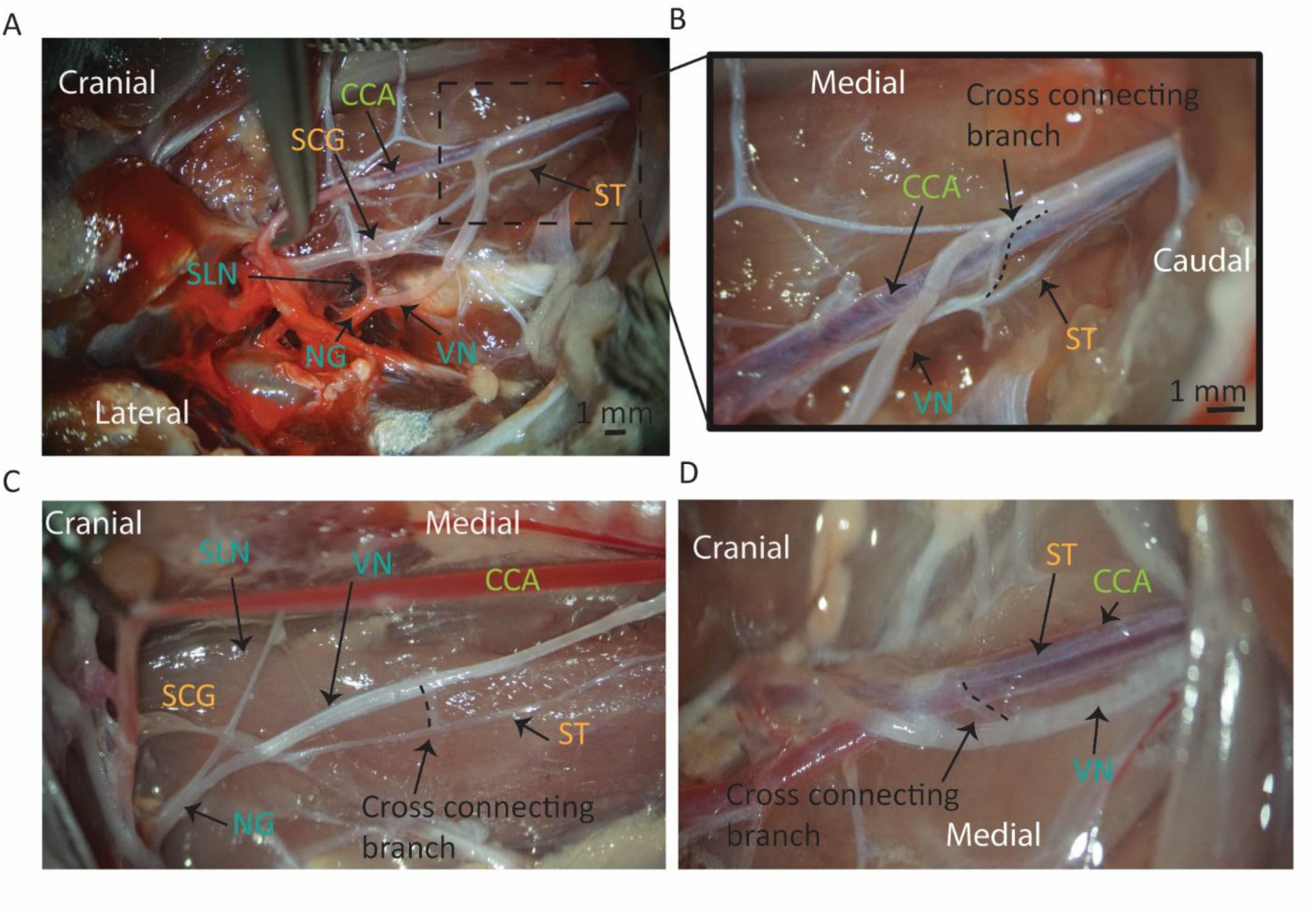
Cross connecting fibers between ST and the VN. A) Anatomical visualization of ST dissected from SCG and cervical VN from NG, with carotid bifurcation retracted to the medial side. SLN originating at the NG and terminating at the neck muscles on the medial side. Cross connecting branch with ST and VN (dotted black box) B) Zoomed in image of dotted black box from panel A shows cross connecting branch between the cervical VN and ST at the common cuff implantation location C) Cross connecting branch from the ST to the VN with the carotid artery retracted medially D) Microdissection of left cervical ST and the VN with cross connecting branch between them near caudal end of the surgical pocket. SLN: Superior laryngeal nerve; SCG: Superior cervical ganglia; NG: Nodose Ganglia; ST: Sympathetic Trunk; VN: Vagus Nerve; CCA: Common carotid artery. Additional pictures of cross connecting branches between the ST and the VN in Supplementary figure 9.

The ST maintains very close proximity with the cervical VN (∼ <2 mm) along the stretch from the SCG to the sternum, located together in a bundle in the carotid sheath (n=10/10 right, n=8/10 left). Importantly, it frequently runs conjoined with or in proximity to the cervical VN, at the most common epineural cuff placement location (dorsal and caudal to omohyoid muscle, See Figure 2B). Consequently, the ST would frequently be included in an epineural cuff placed on the VN at this location (n=9/10 right, n=7/10 left) [Figure 2 and Supplementary Figure 5-7]. There were some exceptions wherein the ST was on the opposite side of the carotid artery than the VN (n=3/10 left) [Supplementary Figure 8]. However, the ST was still part of the carotid bundle, thus having a high likelihood of being cuffed and stimulated with the VN.

### 2) There are cross connecting branches between the cervical VN and ST

In rodents, separating the ST and the VN caudal to the SCG revealed cross connecting branches between the two nerves (n=6 out of 10, right; n=4 out of 10, left). The frequency and location of these branches was highly variable between subjects. One branch separated 1-2 mm caudal to the SCG (4 out 10, right) joining the VN. All branches found were extremely small and fragile within a range of 30-50 µm in diameter.

### 3) Cross connecting branches between VN and ST imaged using MicroCT

We imaged 3 rat cadaver heads using microCT to verify the presence of connecting branches between the VN and ST that may be impacted by an invasive cutdown required for microdissection. This technique ensured that the anatomical area of interest from the carotid bifurcation to the sternum was unperturbed. The SCG was identified on the dorsal side of the carotid bifurcation (n=2), with its size and location comparable to dissection results [Figure 4 B and C]. There was a plexus of numerous nerve branches originating from the SCG traveling to the carotid bifurcation, VN, glossopharyngeal nerve and surrounding muscles [Figure 4C].

**Figure 4:**
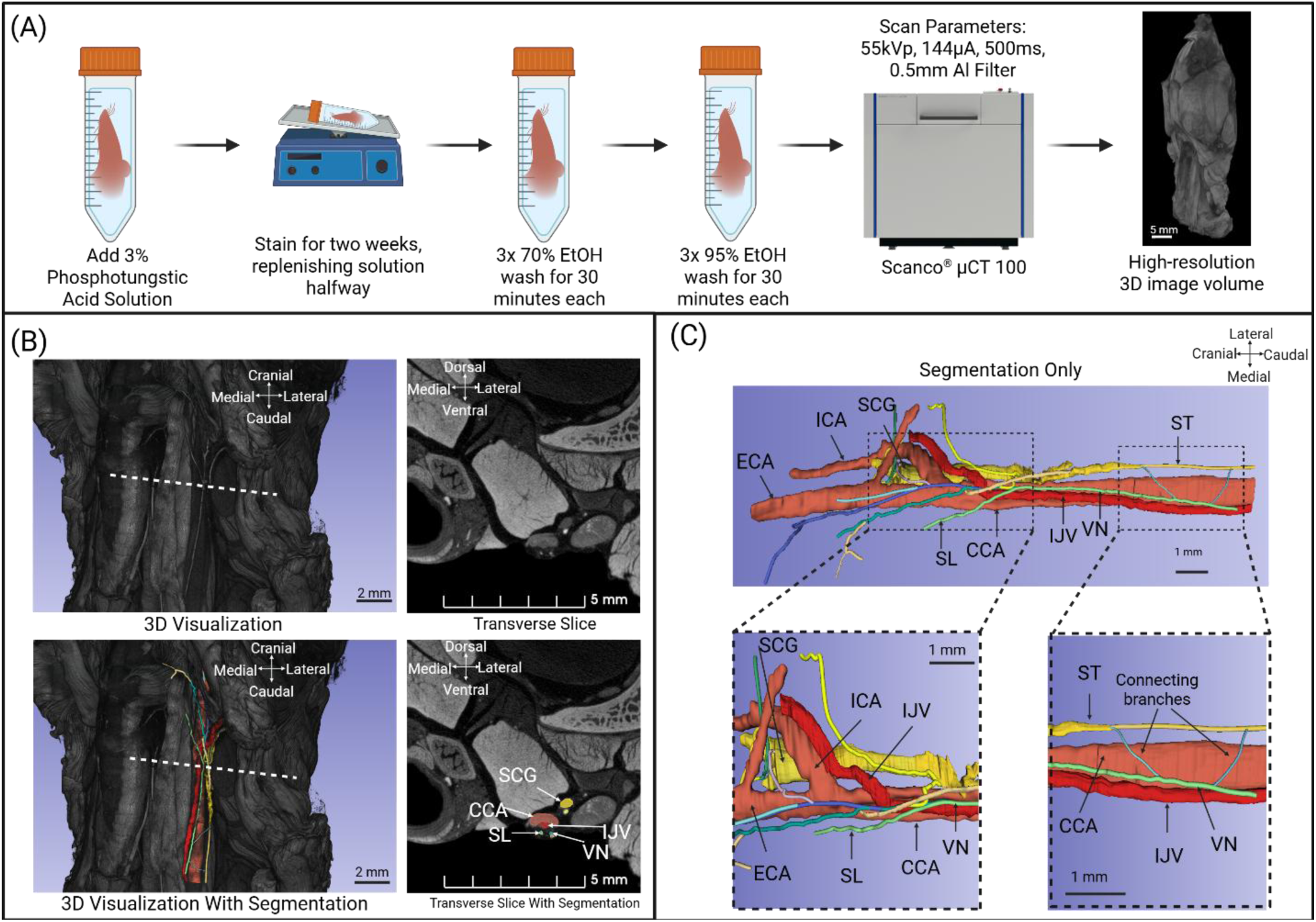
(A) A summary of sample preparation and scanning methodology. (B) A close-up 3D visualization of the left side of the cervical region by the carotid bifurcation alongside a transverse slice along the dashed white line. Manual segmentation was performed to outline nerves along the transverse plane. (C) The segmentations are isolated from the 3D volume. Cross connections between the ST and the VN can be viewed between the VN and ST caudal of the bifurcation. Note: The ST and VN section represented in the figure is ∼6 mm from the carotid bifurcation, hence considerably cranial from the common cuff placement location. We were unable to reliably trace the ST beyond this point due to considerable shrinkage of tissue due to the ethanol baths and formalin preservation(Paramasivan et al., 2021), in addition to the location of the cut at the clavicle. Hence, there is a high possibility that the ST might be joint to the VN at the common cuff placement location (∼9-10 mm caudal to the carotid bifurcation). ICA - internal carotid artery, ECA: external carotid artery, IJV- internal jugular vein, CCA – common carotid artery, SL- superior laryngeal nerve, VN - vagus nerve, SCG – superior cervical ganglia, ST – Sympathetic trunk. This figure was Created in https://BioRender.com.

The ST could not be reliably traced beyond ∼5 mm caudal to the SCG using microCT. This was possibly due to the small size of the rodent ST, inconsistent staining, inadequate contrast and lack of resolution in the microCT imaging. Hence the entire course of the ST (from the SCG to the sternum), especially at common cuff implantation location (∼9-10 mm caudal to the bifurcation) could not be verified with microCT.

Otherwise, the microCT data confirm the results of our microdissection data [Supplementary figure 3 and 4] and previously published literature (Hedger & Webber, 1976; Mitsuoka et al., 2017) and suggest there are additional cross-connections between the VN and ST that are not visible to the eye during microdissection. The ST was identified as the nerve trunk originating from the caudal end of the SCG.

The VN was identified as the other nerve running parallel to the carotid artery. The SLN branched from the VN at the carotid bifurcation and terminated in the muscle group in the neck area (muscle not shown in the figure). The ST and the VN were visualized [Figure 4 B, C] running in parallel along the carotid artery in proximity to each other.

Connecting branches between the ST and VN, 1-2 mm caudal to the carotid bifurcation and SCG were observed using microCT [Figure 4 Panel C]. Multiple other micro branches were imaged between the SCG and VN, cranial to the carotid bifurcation, which were too small to see by eye and may have been cut during microdissections. These findings show the value of microCT imaging as a technique to overcome the limitations of a traditional microdissection cutdown to study the cross-connecting branches between these two nerves.

### 4) Tyrosine hydroxylase (TH) + post ganglionic sympathetic fibers run along and within cervical VN

Rodent cervical carotid bundle cross sections were stained with TH+ to analyze the hitchhiking ST with the VN (entire dataset available on SPARC website (Pelot et al., 2022); a subset is reported and analyzed here [n=6 sourced from Duke University] with author permission). Locations of these cross sections within the neck varied and could not be reliably matched to the location where the ST and VN were conjoined during our microdissections.

The samples contained TH+ fibers bundled together surrounded by perineurium in the carotid sheath. This nerve was annotated as the ST (Figure 5 A-F) with its average diameter being 89.22±24.03 um.

**Figure 5:**
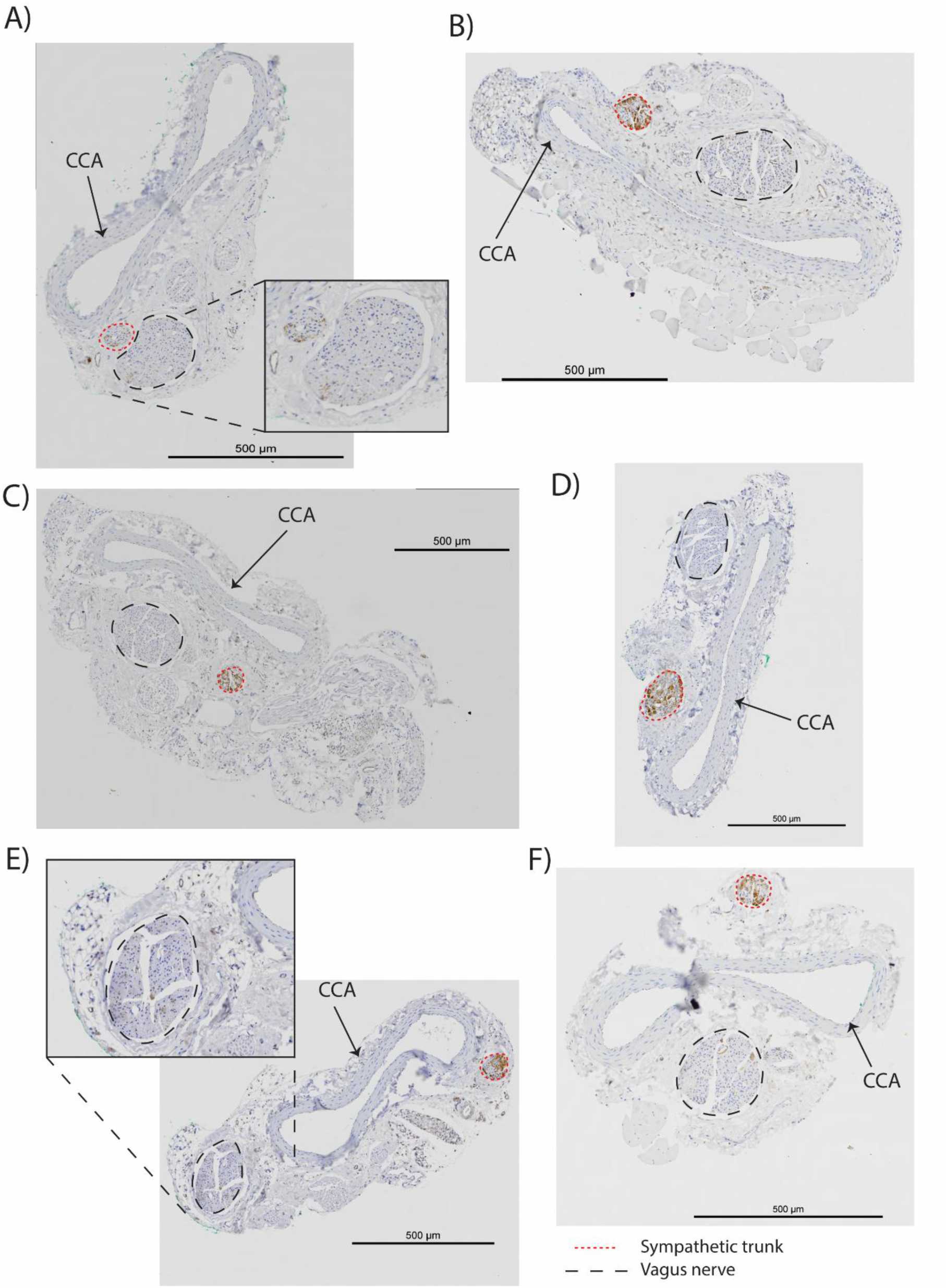
Tyrosine hydroxylase (TH+) post ganglionic sympathetic fibers run along and within the VN. Cervical carotid sheath samples were fixed and stained with Tyrosine hydroxylase (TH+) to identify post-ganglionic sympathetic fibers. Grouping of TH+ fibers in a fascicle identified as ST (red dotted circle) which runs with the cervical VN (black dotted circle) in the carotid sheath. **A-D)** The TH+ ST fibers were found in proximity or running close to VN bundle on the same side of the carotid artery (purple smooth muscle arterial wall). **E-F)** ST was on the opposite side of the VN, in reference to the carotid artery in a subset of samples. The TH+ positive fibers were also found in the VN bundle itself **(Panel A and E inset box)** independent of ST bundle. The entire dataset can be found at <u>Rat VN TH- (tyrosine hydroxylase) and</u> <u>ChAT- (choline acetyltransferase) positive fibers (sparc.science)</u>.

Another larger grouping of fascicles surrounded by perineurium running parallel to the carotid artery was identified as the VN. The majority of the fibers from this bundle were TH-. The ST was “hitchhiking” with the VN (average edge to edge distance of 221.25 ± 141.08 um between the nerves, individual distances in Supplementary table 1) on the same side of the carotid artery (66.67% of the samples) (Figure 5 panel A-D). In 33.33% of samples, the ST was found on the opposite side of the VN, with the carotid artery in the middle (Figure 5 Panel E and F). In addition to the ST bundle, TH+ fibers were also observed in the VN itself (Figure 5 A, B, E and F). Other nerves were also present in the carotid sheath, which were comparable in diameter to the ST but not TH+. These additional nerves could be the aortic depressor nerve (ADN) or small branches of VN itself (Figure 5 A, B, C and E).

These histological results in conjunction with anatomical data support the hypothesis that the ST hitchhikes alongside the VN within the carotid sheath at the cervical level for cuff placement.

### 5) Functional implications of stimulating hitchhiking ST during VNS

Increase in bradycardic response (Mean ± std dev) to increasing VNS amplitude (0-500 µA) across the cohort G) Increase in tachycardic response (Mean ± std dev) to increasing STS intensity (100-800 µA) across cohort H) Loss of VNS evoked bradycardic response post VN transection caudal to stimulation electrodes I) Loss of STS tachycardiac response post ST transection caudal to stimulation electrodes. All data in the figure is from the right side. Similar responses were recorded from the left side, see Supplementary Figure 10.

The ST and VN were stimulated independently with increasing stimulation amplitudes, to record the corresponding changes in HR (n=8 right, n=3 left). The stimulation location was selected such as the two nerves could be interfaced independently, but as close to the most widely reported cuff location (caudal or dorsal to the omohyoid muscle) where they are conjoined. In cases where the two nerves were conjoint ∼2-3mm from the carotid bifurcation, the two nerves were carefully separated to create enough space to interface the two electrodes while avoiding visible cross connections between them.

A VNS dose dependent increase in bradycardic response was recorded in all subjects (n=8, right, n=3 left) [Figure 6 D, Supplementary Figure 10 panel B, D, F]. The maximum drop (bradycardia) recorded was 100 beats per minute [BPM]. Stimulation of isolated VN evoked bradycardia, which was immediately followed by a clear tachycardiac rebound response in all subjects. Stimulating the isolated ST caused a consistent tachycardic response (n=7 right, n=3 left), with increased stimulation amplitude causing a greater increase in HR [Figure 6 E, Supplementary Figure 10 panel A, C, E]. The thresholds for causing VNS-induced bradycardia were lower (20-50uA) compared to STS caused tachycardia (100-200uA).

**Figure 6:**
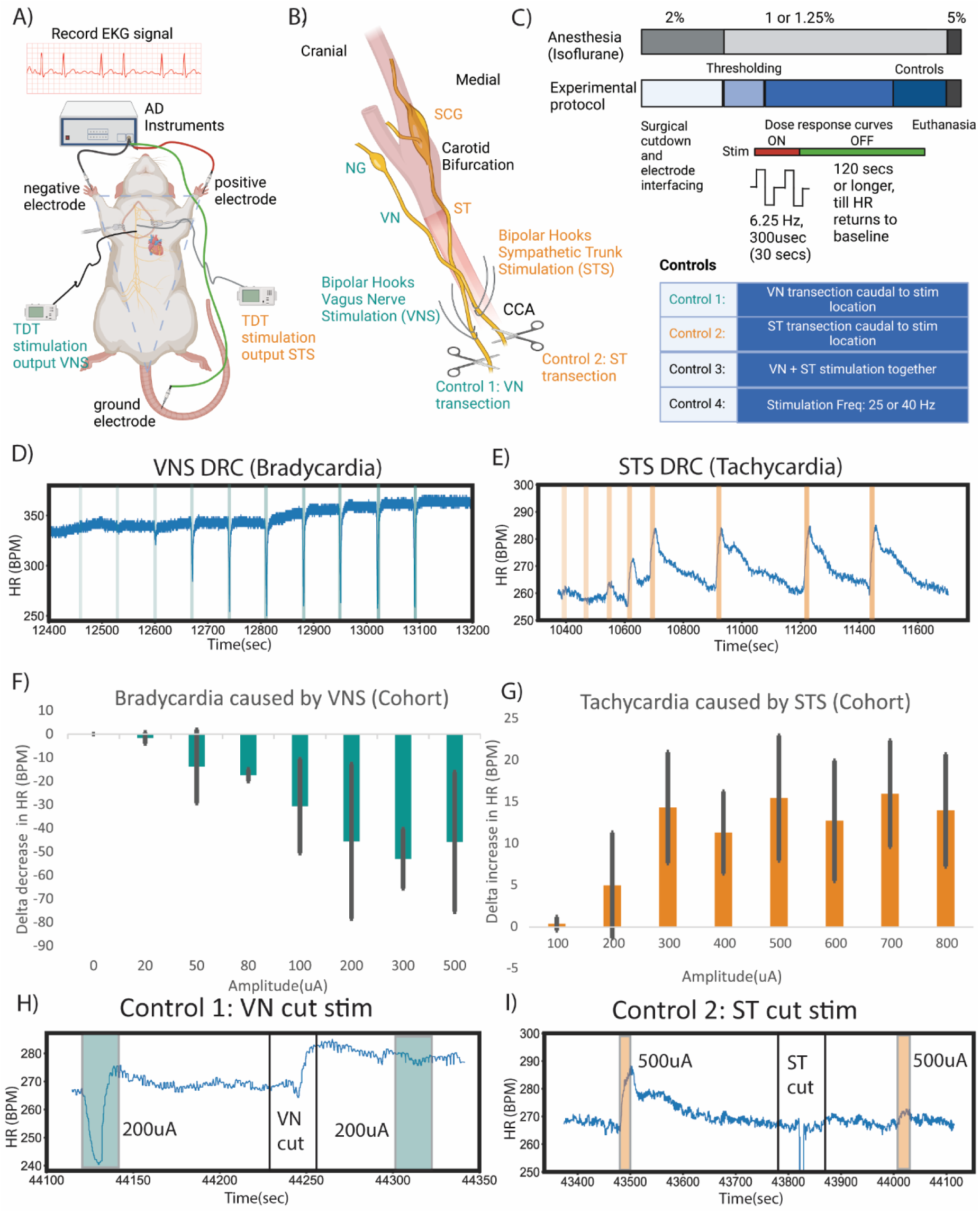
Effects of vagus nerve stimulation (VNS) and sympathetic trunk stimulation (STS) on heart rate: Illustrative drawing of the experimental setup A) Cardiograms recorded using a 3-electrode set-up using AD Instruments and two independent TDT current sources for VNS and STS B) Two pairs of bipolar hook electrodes for VNS and STS at common cuff implantation location. Transection of nerves caudal to stimulation location as controls for on-target engagement. C) Experimental timeline: Isoflurane level was 2% during sedation/surgical cutdown and then lowered to 1-1.25 % during collection of functional data. Stimulation was delivered at 6.25 Hz with 300 µsec pulse width (150 µsec per phase) with 25 Hz and 40 Hz as additional controls. Dose response curves were collected by stimulating at, below and above threshold amplitudes with ON time of 30 secs and OFF time of 2 mins or longer (till baseline returned to pre-stimulation value) D) VNS dose dependent bradycardia in HR (beats per minute) from one representative animal. Highlighted blue bands represent stimulation with increasing stimulation amplitude (0-800 µA) E) STS dose dependent tachycardiac response in HR (beats per minute) from one representative animal. Highlighted orange bands represent the stimulation timing (100-800 µA) F)

There was a clear overlap between the two dose response curves (200-500uA) [Figure 6 F, G], wherein these amplitudes caused bradycardia during VNS and clear tachycardia during ST stimulation. For example, at 500 µA (a stimulation amplitude repeated in n=6 animals, right side) the average VNS bradycardic response was 23.16 ± 63.06 BPM and STS tachycardic response was 16.08 ± 8.12 BPM. Interestingly, this overlap amplitude range is in the rodent therapeutic window for various VNS applications(Manta et al., 2009). The VNS-induced drop reached maximum change and returned to baseline instantaneously whereas there was a delay for STS-induced tachycardia to reach its peak delta change and its subsequent return to baseline [Figure 6 D, E, H, I and Supplementary Figure 11]. This points to possible different mechanisms and their corresponding long-term effects on baseline HR.

To verify stimulation driven bradycardia and tachycardia were due activating fibers in the VN and ST respectively (on-target engagement), stimulation was applied before and after the main nerve trunks were sequentially transected. The nerve transections were caudal to the stimulation electrodes while keeping all other nerve branches intact. This would putatively help distinguish between efferent and afferent nerve pathways for cause and effect. There were no stimulation evoked effects on HR post nerve transections on the left and right side. Paired data of delta HR changes for above threshold stimulation amplitudes pre and post transections was analyzed (n=4 animals). VNS and STS HR changes were significantly reduced after VN and ST transections, respectively (Paired one tail t-test, normality verified using Shapiro-Wilk test, VN p=0.023, STS p = -.002) [Figure 6 H, I and Supplementary Figure 10 E, F, Supplementary Figure 14 and 15].

Previous studies have reported that stimulating the conjoint ST and the VN results in attenuated bradycardic response (∼60% reduction in baseline HR) as compared to VNS alone (∼85% reduction in baseline HR)(Randall et al., 1969; Yang & Levy, 1992) in canines. In a rodent study, background stimulation of the isolated ST (5 Hz) with the VN (25,40 Hz) was reported to have an antagonist effect towards VNS on HR (Kawada et al., 2023). To recreate the ST accidentally being cuffed along with the VN during rodent VNS studies, the ST and the VN were stimulated in conjunction with the same stimulation amplitude as a pilot study. This was implemented using two methods, the first, with one hook electrode wrapped around the conjoint nerves and the second, stimulating the separated nerves with the same amplitude at the same time. Both methods yielded bradycardia at the stimulation parameters tested of 100, 200 and 800 microamps (n=4 animals) [Supplementary Figure 16 and Supplementary Figure 17]. However, at these stimulation parameters VNS+STS did not elicit a statistically significant attenuated bradycardic response as compared to VNS alone given the small number of animals and the lack of granularity of the dosing (Wilcoxon-Signed Rank test, Z= -1.578, p = 0.115, N=17 paired observations for n=4 animals). This pilot study was only powered to detect an effect size of 0.383 (r=Z/√N). Given 1) the threshold for bradycardia was 20-50 microamps with VNS, 2) the amount of current our electrode was delivering to the ST and VN individually when cuffed together is unclear and would be very specific to our electrode design and 3) the number of animals required to be appropriately powered to detect a small difference in degree of bradycardia, it was not deemed ethical, practical or meaningful to attempt to increase the number of animals to detect a smaller mean difference.

All the above reported data is for stimulation frequency of 6.25 Hz. However, additional clinically relevant frequencies of 25 Hz and 40 Hz (n=2 right, n=2 left) were tested in a subset of subjects. STS and VNS respective tachycardic and bradycardic responses were also reliably observed at 25 and 40 Hz [Supplementary Figure 18].

## Discussion

In this paper, we establish that the ST is frequently conjoined with the right and left VN at a common location for VNS in rodents. Using microCT, we also demonstrate there are cross-connections between the ST and the VN similar to what has been reported in humans that are difficult to identify by eye in acute or chronic VNS surgical window. Stimulating the ST between the MCG and SCG where it is isolated from the VN reliably causes tachycardia starting at ∼200 microamps, well within the range commonly used for rodent VNS studies. These data in aggregate suggest that the contribution of sympathetic fibers to the stimulation response of VNS in rodents, as well as unintended surgical dissection of these cross-connecting pathways, should be considered in mechanistic studies of VNS. In the following discussion, we outline the implication of these results for multiple VNS therapies, as well as best practices for identifying ST contributions to VNS.

### Potential implications of ST as an off-target nerve in VNS functional studies

#### Interpretation of biomarkers for VNS on-target engagement

There have been several efforts to develop biomarkers to show on-target engagement of specific fiber types during VNS (Blanz et al., 2022; Chang et al., 2020; Gurel et al., 2020). VNS-induced bradycardia is an established biomarker for activation of smaller diameter efferent parasympathetic b-fibers (Higgins et al., 1973; Löffelholz & Pappano, 1985; Qing et al., 2018). Bradycardia is frequently used to confirm the viability of electrode-nerve interface in long term chronic rodent studies. Additionally, stimulation parameters are often titrated until bradycardia (∼5-10 BPM) is recorded and then reduced by 70-90% of that value to target activation of vagal sensory fibers (Huffman et al., 2019).

This paper reports that the ST can hitchhike with the VN at the traditional cuff implantation location and causes bradycardia when stimulated together. Thus, reporting evoked bradycardia or lack of tachycardia as a biomarker for not activating the ST, is insufficient in rodents. Additionally, concurrent stimulation of both the ST and the VN can lead to highly variable bradycardia thresholds compared to stimulating VN alone; due to accentuated antagonism and reciprocal excitation (Kawada et al., 2019; Yang & Levy, 1992). Thus, it would result in highly variable stimulation amplitudes in individual animals, potentially leading to variable dosing results across a cohort.

Another biomarker used to show afferent VNS target engagement is pupillometry. The cranial efferent postganglionic branches from the SCG are known to innervate the pupil and cause dilation; with sympathetic input controlling the dilator muscles and the vagal parasympathetic output controlling sphincter muscles (Bianca & Komisaruk, 2007; Liu et al., 2017). This has led studies to use pupillometry as a biomarker for activating afferent VN fibers and to parameterize therapy (Mridha et al., 2021).

However, if the ST fibers are stimulated along with the VN, their input to the SCG may also drive the dilation of the pupil. Moreover, transections cranial to the cuff are often used to verify VNS induced pupil changes are caused by activation of vagal afferents. As the ST and VN are frequently conjoined at the most common electrode cuff location, a cut cranial to the electrode would sever both stimulated vagus afferent input to the brain as well as pre-ganglionic sympathetic projections to the SCG.

It is also important to note that many studies demonstrate off-target muscle activation during VNS through multiple nerve pathways, including the recurrent and superior laryngeal pathways. Any noxious or ‘surprising’ sensation may also change the arousal and therefore pupil diameter. Given the multitude of motor nerves in close proximity to the cervical vagus, it may be functionally difficult to selectively transect ‘just the cervical vagus’ with precision.

#### Variable efficacy in VNS dose response studies

Preclinical rodent models of VNS are used as a starting point for mechanistic studies for epilepsy (Handforth & Krahl, 2001), inflammation (Carvalho et al., 2013; Olofsson et al., 2015), depression (Biggio et al., 2009; Manta et al., 2013; Revesz et al., 2008), heart failure (M. Li et al., 2004) and others. Importantly, the known pathophysiology of many of these indications include higher than normal sympathetic signaling to the peripheral organs (Hausberg et al., 2007; Meltzer & Meltzer, 1903; Pongratz & Straub, 2014), suggesting an altered autonomic balance biased towards sympathetic overdrive. As we demonstrated in this study, there was a clear overlap in rodent VNS therapeutic window amplitudes (200 µA-800 µA) where both nerves could be activated.

Additionally, some VNS rodent studies report a reduction in therapy efficacy (inverted U- dose response curve) in the stimulation amplitude range of 500-1500 µA (Manta et al., 2009; Pruitt et al., 2021; Souza et al., 2021; Zuo et al., 2007). In our experimental data, we reached a saturated ST activation response in similar stimulation amplitude range [Figure 6 G]. Accidental activation of the ST at higher doses could be a contributing factor, in addition to reported varying neuronal population firing in the NTS, for mixed VNS experimental results unless isolating the VN from the ST is clearly stated in the surgical protocol (Clark et al., 1995).

### Accidental activation of the sympathetic system could worsen disease prognosis

In most chronic diseased conditions, sympathetic activity is persistently elevated and can trigger long term detrimental pathological remodeling(Elefteriou et al., 2014; Gardner et al., 2016; Shi et al., 2014). In this paper we demonstrated stimulation parameters as low as 200 microAmps may activate sympathetic fibers in rodents, which may exacerbate underlying pathology. For example in studies using VNS to treat heart failure, simultaneous stimulation of the parasympathetic and sympathetic systems may increase sympathetic outflow to the heart in addition to increasing parasympathetic outflow; increased sympathetic outflow has been shown to increase the risk of atrial fibrillation (Sheng et al., 2011). These data suggest it may be critical to avoid activating the sympathetic system during VNS clinical studies to help improve patient outcomes, which may have contributed to several recent failed VNS clinical trials in heart failure (Anand et al., 2020). Similarly, in VNS studies intended to activate the inflammatory reflex, partial activation of the sympathetic system could lead to different levels of inflammatory cytokines expressions as compared to activation of the parasympathetic system alone. While the effector arm of the sympathetic system is organized along different spinal levels (Jänig & McLachlan, 1992), previous literature has reported the effects of cervical sympathetic components on peripheral organ functions at other spinal levels (Ahlman et al., 1978; Cardinali et al., 1981; Zhang et al., 2022).

### Histology Considerations and Recommendations for Best Surgical Practices

#### Histological studies of the ST and its connections

The sympathetic fibers, in the ST and in the VN itself, were verified by histological data (TH+ staining). TH+ post ganglionic sympathetic fibers would mostly be small diameter unmyelinated c-fibers (Keast, 1995b) and would have a higher threshold of activation than parasympathetic b-fibers, according to conventional fiber diameter recruitment curves (Erlanger & Gasser, 1930). However, preganglionic cholinergic sympathetic fibers would also be present in the cervical ST, with similar size and threshold as parasympathetic efferent b-fibers (C. Li & Horn, 2006).

Since TH+ stains exclusively for post-ganglionic sympathetic fibers and not pre-ganglionic sympathetic fibers (ascending in the sympathetic trunk from Middle Cervical Ganglia to SCG (Asamoto, 2005; Billingsley & Ranson, 1918; Foley, 1943)), there could be a much higher number of sympathetic fibers in the VN than reported. Future work should include multiple IHC stains such as CHAT+ (Kronsteiner et al., 2024b) along with tracing to the SCG, to help elucidate all different sympathetic fiber types and their organization in the cervical VN. Future work should focus on techniques for assessing the role of these pre-ganglionic sympathetic fibers in the sympathetic trunk and cross-connections with the VN. It could also help with targeting specific cervical sympathetic connections to the lymph nodes, thyroid glands etc. to study the effect of sympathetic signaling on their functions.

### Surgical cutdown and electrode placement for cervical VN in rodent studies

In our initial review of the rodent VNS literature, the potential mechanistic confounds of the ST conjoined with the vagus, or cross connections between the ST and VN, were largely overlooked. A common unstated assumption was that ST nerve fibers are not located within the carotid sheathe, and therefore any nerve cuffed with an electrode that resides within the sheathe will contain only vagal fibers. Contrary to this assumption, we report the presence of the ST in the carotid sheath, “hitchhiking” with the VN for long stretches along the cervical region in the rodent model. This was highly likely at the traditional cuff implantation locations (placed cranial or caudal to the omohyoid muscle), putatively making the ST a potential source of off-target effects during mechanistic studies of VNS.

An improved surgical protocol, with additional steps to identify the ST could help avoid potential confounds. As the aortic depressor nerve (ADN) can also sometimes run with the VN in the carotid sheath (Krieger & Marseillan, 1963) and is comparable in size to the ST, identifying and tracing the ST from the SCG is the most reliable technique. In chronic studies, where the surgical window is limited, the ST can still be identified as the smaller diameter nerve running in parallel to the VN next to the carotid artery. A quick test stimulation of the small nerve bundle can be used to distinguish between the ST and the ADN as stimulating the ST causes tachycardia while stimulating the ADN causes bradycardia (Pinto et al., 2016). Additionally at study termination, postmortem dissection studies should be performed to ensure the ST was not accidentally cuffed, by tracing it back to the SCG. For clinical studies, care must be taken during isolating hitchhiking ST from the VN, as damage to the ST and SCG can lead to Horner’s syndrome which is a reported side effect in VNS clinical trials (Cozzaglio et al., 2008; Genovese et al., 2020).

#### Cross connecting branches between the ST and the VN

This paper demonstrated there are frequently connecting branches between the ST and the cervical VN in rodents. It is possible that cross-connecting branches were present in more subjects than reported but were either damaged during microdissection or remained undiscovered due to their extremely small diameters. The variation in the anatomical location and frequency of these branches along the cranial-caudal axis, points to a need to be cautious of their presence during microdissection. If severing the branches between the two nerves cannot be avoided, it could be a possible limitation in the experimental design.

Recently published articles documented these cross connections in humans (Franco-Riveros et al., 2024; Seki et al., 2014), canine (Chase & Ranson, 1914) and swine (Settell et al., 2020). These data in aggregate suggest that frequent micro scale cross connections between the ST and VN beyond the canonical large branches is a cross-species phenomenon, possibly highlighting it as a consideration in translational research. Human fetal cadaver studies report these branches between the ST and VN in the cervical region can coalesce and form the cardiac nerves (González et al., 2023). As our experiments were not designed to study the physiological implications of these branches in cardiac neuroaxis or towards other indications, the implication towards VNS clinical translation for cardiac control, inflammation remains to be explored. These branches may suggest a local cross-communication system between the two autonomic systems working synchronously to maintain homeostasis, perhaps independent of higher order processing in the cortex.

### Recommendation for VNS surgical cutdown to improve experimental methods for rodent VNS studies

1. Carefully identify the ST (∼1/8th size of the VN) present in the carotid sheath.

a. Identify the neural bundle in the carotid sheath.
b. Observe the neural bundle (under a surgical microscope) to visualize if a smaller nerve (∼200 µm diameter) is running conjoint with the VN (∼800 µm diameter)
c. Verify if it is the ST by tracing it cranially to the SCG.
d. If the surgical window is limited during chronic electrode implantation, make a note if the ST is conjoint to the VN before electrode implantation. Try to implant electrode slightly caudal/cranial to this location if the two nerves are found running separately.
2. If the ST is found joint to the VN in the carotid sheath during electrode placement, try to gently blunt dissect the ST from the VN using fine forceps or glass pipette rods to prevent stimulating both nerves.
3. Be aware of cross connections between the VN and ST while separating the two nerves.
4. If separating the ST from the VN is not feasible and would likely be stimulated, consider it as a possible caveat in VNS mechanistic studies or address using additional controls. Confirm and report if the ST was cuffed with the VN, with post-mortem dissections and histology at the termination of the study.

Given the ST also hitchhikes in large animals and humans, understanding the involvement of activation of the ST in potential effects and side effects is critical for mechanistic understanding and clinical relevance of animal studies. Studies need to clearly report what they intended to stimulate (isolated VN or the vagosympathetic trunk), how they surgically interfaced with it and functionally confirmed the effects of their stimulation with appropriate biomarkers. Without this information it is more difficult to appropri- ately assess fundamental mechanisms of actions or how outcomes will translate to the clinic. Moreover, comparisons/replication across studies becomes problematic.

## Overall Study Limitations

There are several limitations of this paper which have been broken into major categories below.

### Microdissections

The microdissection studies were not statistically powered to objectively capture all the anatomical variations of the ST regarding left and right branching patterns, or gender differences.

This study was not designed to cover all variances in cross connecting branches between the ST and the VN and should be treated as an observation and an exploratory result. The low number of subjects, with the possibility of cutting smaller branches during dissections, limits this study from investigating the complexity of size and distributions of the cross-connecting branches between the two nerves.

### TH+ Histological staining

The TH+ staining was conducted in a different cohort of subjects and was not matched with anatomical dissection or functional data. The samples were collected as a carotid bundle (ST+VN+Carotid artery) with the carotid sheathe intact. However, the location of the cross section along the cranial-caudal access was not annotated. Hence, there is a high likelihood that the ST and the VN were conjoint in locations not captured in the histology sections reported in the paper. Future rodent studies should have serial histology tracing from the SCG to the sternum, as previously reported in humans (Ruigrok et al., 2023). Additionally, while the location of the ST with reference to the VN and the carotid artery was reported at the location where the sample was collected, it was not possible to annotate if the ST was medial/lateral to other anatomical landmarks.

### MicroCT data

MicroCT imaging limitations included variable stain uptake which causes dark regions deep in the tissue. This variable staining led to some neural features being difficult to visualize as compared to surrounding muscle layers. While the nerves should have minimal dislocation as compared to microdissected samples, disarticulation of the head above the clavicle could have caused loss of tension in the nerves and hence some movement. Shrinkage of the sample tissue due to chemical fixatives should also be noted while comparing nerve sizes to microdissection data.

### Functional data

This paper explores if the ST can be activated during VNS rodent studies and its functional effects on HR variability. All the functional data was collected under anesthesia, which is known to alter the stimulation thresholds. Most of the functional data was collected using a stimulation frequency of 6.25 Hz, which is lower than the frequencies used for clinical applications (eg. 25 Hz). While our pilot experiments using 25 Hz also showed an overlap for activation thresholds between the two nerves, additional future studies are required to completely map out the stimulation dose response curves at relevant frequencies. The stimulation amplitudes were not randomized while acquiring the HR changes. Hence, there could be additive effects of increasing stimulation amplitude on the HR responses. Historically the therapeutic target for studying heart failure is the right VN; hence our cohort is heavily biased towards right VNS (n=8). However, since left side VNS is used in multiple other mechanistic studies such as epilepsy, inflammatory responses, stroke rehabilitation, etc., a smaller cohort (n = 3) was also added for left VNS.

The experiments were not statistically powered to address the differences in left versus right side stimulation on HR variability. This has been previously reported in other publications (Shao et al., 2021) and are not reported in the paper. Stimulation of the VN and the ST simultaneously yielded bradycardia in our studies . This was different from translationally relevant VNS swine studies (Blanz et al., 2022), that used smaller contact circumferential electrodes to selectivity activate the embedded ST alone. However there were several limitations to duplicate these results in rodents; reliable concurrent current source stimulation (equal charge density delivered to both nerves), lack of good contact of hook electrodes with the ST embedded with the VN (due to size of ST being ∼1/8th of the VN), 270 degree circumferential electrode nerve contact (lack of spatial selectivity), rodent nerves being considerable smaller in size (less separation of distance between fascicles from the stimulation electrode, so entire nerve gets activated quickly). These results also do not explore the implications of ST activation in different VNS mechanistic studies. The functional studies were not preregistered; hence all results reported in the paper should be deemed exploratory with confirmatory studies required in the future.

## Conclusion

These studies demonstrate that the ST frequently hitchhikes with the cervical VN at the common cuff implantation location in a rodent model. This was verified by anatomical microdissection, imaging and histological data. We also annotated fine cross-connecting branches between the two autonomic nerves in this location. VNS evoked consistent bradycardia while ST stimulation evoked tachycardia across the functional cohort, with a clear overlap in the activation window for the two nerves. These data also suggest possible confounds in interpreting HR (bradycardia) as a biomarker for VNS target engagement alone. We hope these results will help new researchers design VNS rodent protocols to avoid (or at least limit) engaging the sympathetic system, aiding in mechanistic studies for epilepsy, depression, stroke plasticity, inflammation, heart failure, and more.

## Acknowledgements

We sincerely thank Dr. Megan Settell, Maria LaLuzerne, Dr. Nikki Pelot, Dr. Julie Savage and Dr. Aaron Suminski for reviewing the manuscript and providing insightful feedback.

## Data Availability

All data that support the findings of this study are included within the article (and any supplementary files). Any additional data are available from the corresponding author upon reasonable request.

