## Supplemental material for "Prevalence of sympathetic fibers within the rat cervical vagus, and functional consequence on physiological effects mediated by vagus nerve stimulation (VNS)"

Supplementary Material:

**Methods:**

1) Electrochemical Impedance Spectroscopy (EIS) for checking quality of electrodes

The impedance of the electrodes was measured using a 3-electrode setup (Silver/silver chloride – Reference electrode, BASi – MF2056; Platinum sheet - Counter electrode, Metroohm with surface area of approximately 1 cm^2^) by running electrode impedance spectroscopy (EIS) using an AUTOLAB potentiostat (model: PGSTAT12). The electrodes were deemed fit for experiment if their impedance was less than 5 k ohm at 1 kHz

2) TH+ staining protocol for VN samples

For histological studies, microdissection of adult Sprague-Dawley rats [n = 6 female, n = 6 male] was carried out and the left cervical VNs were collected. The tissue samples were then placed in 4% (paraformaldehyde) for 2 to 8 days for fixation. The tissues were then embedded using paraffin and serially sectioned to collect 5 um slices. Next, they were stained using an anti-TH antibody (1:250, Abcam, ab112, RRID:AB_297840) as a primary antibody and iotinylated SP-conjugated Affinipure goat anti-rabbit IgG (H+L), 1:500, Jackson ImmunoResearch, 111-065-144, RRID:AB_2337965 as a secondary antibody. The slices were then imaged using a Nikon Ti2 inverted microscope with a DS-Ri2 color CMOS camera (Nikon Instruments Inc.) at 20x to identify TH+ fibers. (Additional details about the protocol can be found at [SPARC_Duke_Grill_OT2-OD025340_VagusNerve_IHC_TH (protocols.io)](https://www.protocols.io/view/sparc-duke-grill-ot2-od025340-vagusnerve-ihc-th-e6nvw6w4dgmk/v2)

3) Protocol used for making glass rods used in microdissection

For handling fragile and small nerve connections and avoiding damage, micropipette glass rod microdissection tools were created using standard aspirating glass pipettes used in invitro cell cultures. The pipette was taped on a stable surface with the small open bore hole accessible. The pipette tip was held on an open flame (small lighter) and melting it’s bore. These tips were then, gently and quickly, molded using the force-pull method to shape them in the desired shape. These tools with curved angles, help with dissecting the finer structures around the superior cervical ganglia (SCG).

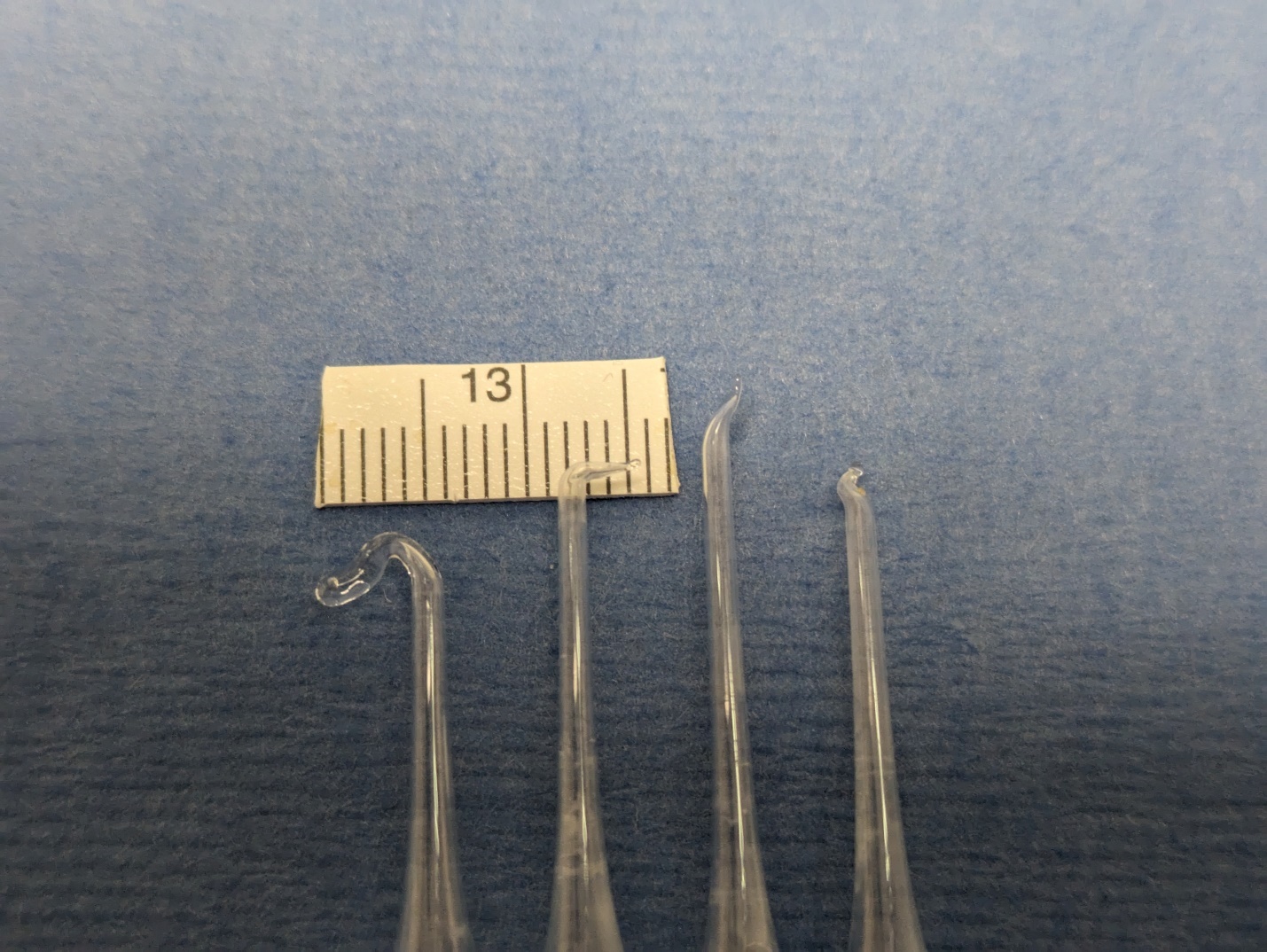

Supplementary Figure 1: Examples of a subset of glass rod microdissection tools. The variation in curvature and sharpness between tools gives flexibility to have a desired surgical approach with different angles in a complex small surgical space at varying depths. Glass material reduces the risk of damage to tissues and bleeding caused by very fine sharp metal instruments. From left to right: Glass rod hook or loop that were used to hold the carotid artery and the VNs, a 90-degree bent retraction tip was commonly used to retract the carotid bifurcation and access the SCG on the dorsal side of the artery. The last two angled tips were used to separate small fibers which were hard to access using conventional fine forceps.

**Results:**

1) *Accessing the superior cervical ganglia and its branches*

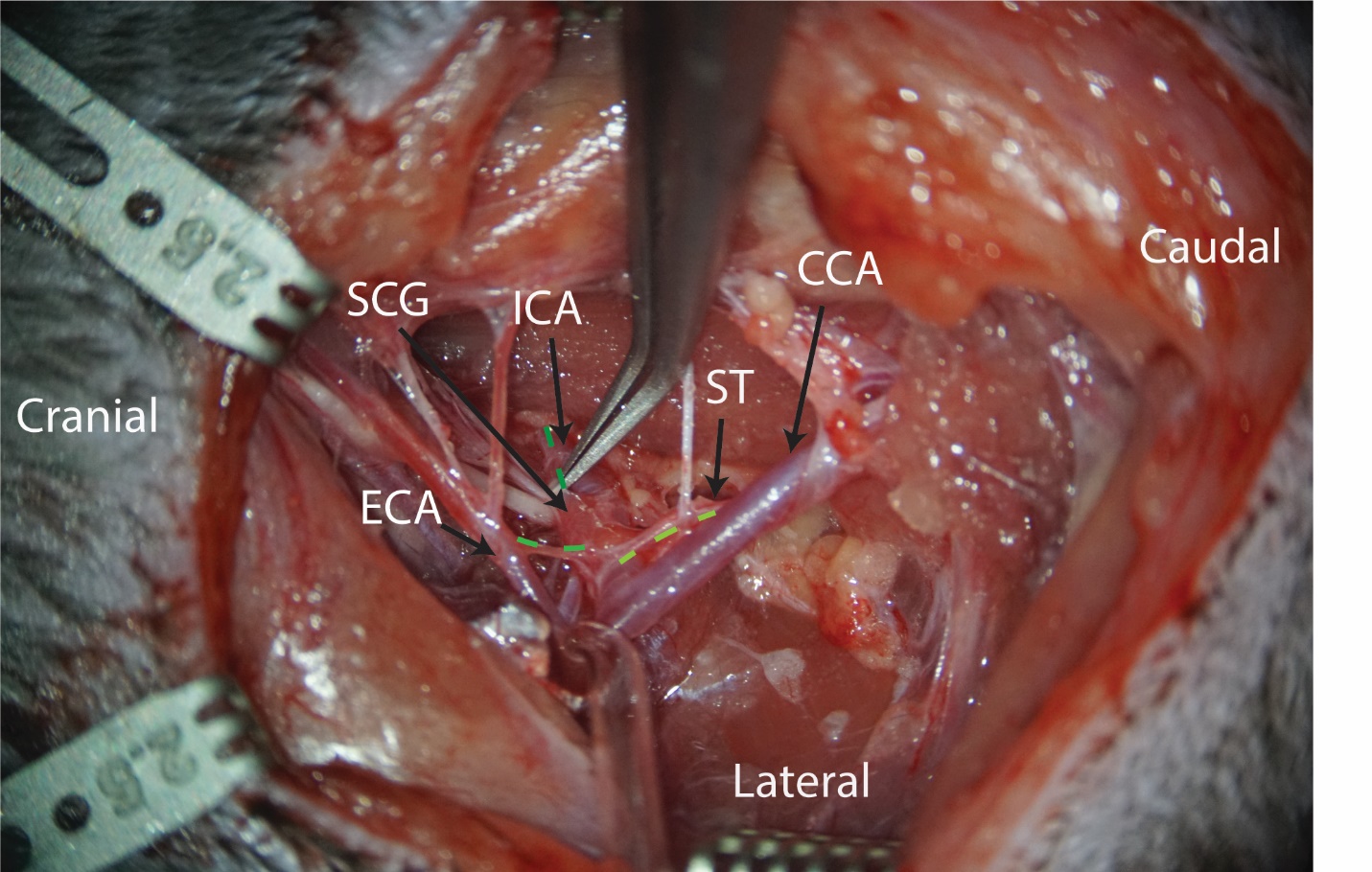

Supplementary Figure 2: **Superior cervical ganglia (SCG) location on dorsal side of carotid bifurcation**: SCG was located dorsal to the carotid bifurcation “sticking” to the interval carotid artery (ICA) with a branch from SCG traveling along the ICA, and a branch connecting external carotid artery (ECA) and ST (ST) originating at the caudal pole end of SCG and traveling caudal along common carotid artery (CCA). (All branches of SCG denoted in green dotted line. The lines are darker to lighter as go from dorsal to ventral).

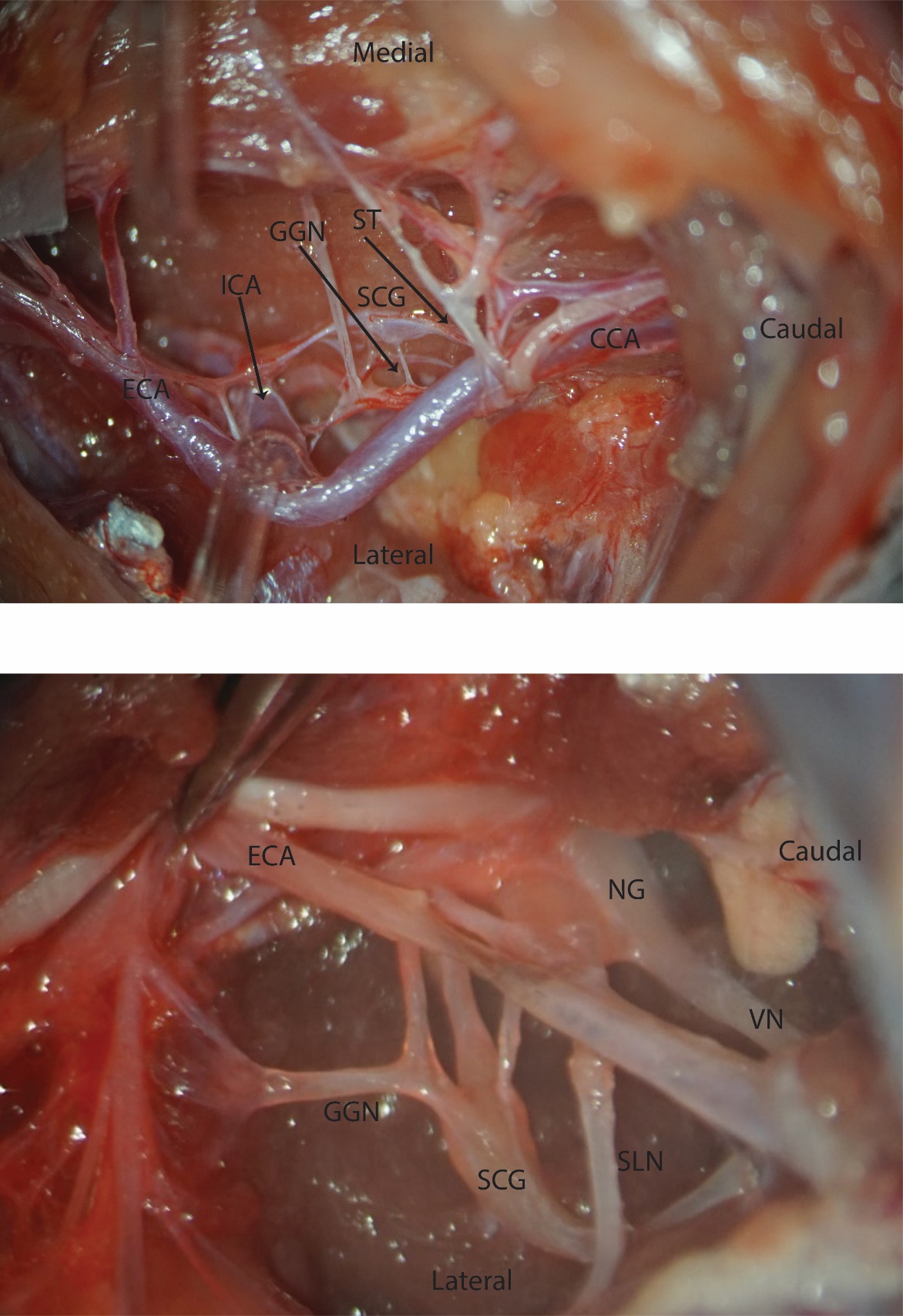

Supplementary Figure 3: **Superior cervical ganglia (SCG) location and identification of its branches.** The SCG had multiple branches which were found following or joining internal carotid artery. The ganglioglomerular nerve (GGN) joined the carotid body, external carotid artery, nodose ganglia. The ST(ST) originated at the caudal pole of the ganglia and traveled along VN and common carotid artery.

2) *Additional cranial nerves in the surgical pocket which were kept intact*

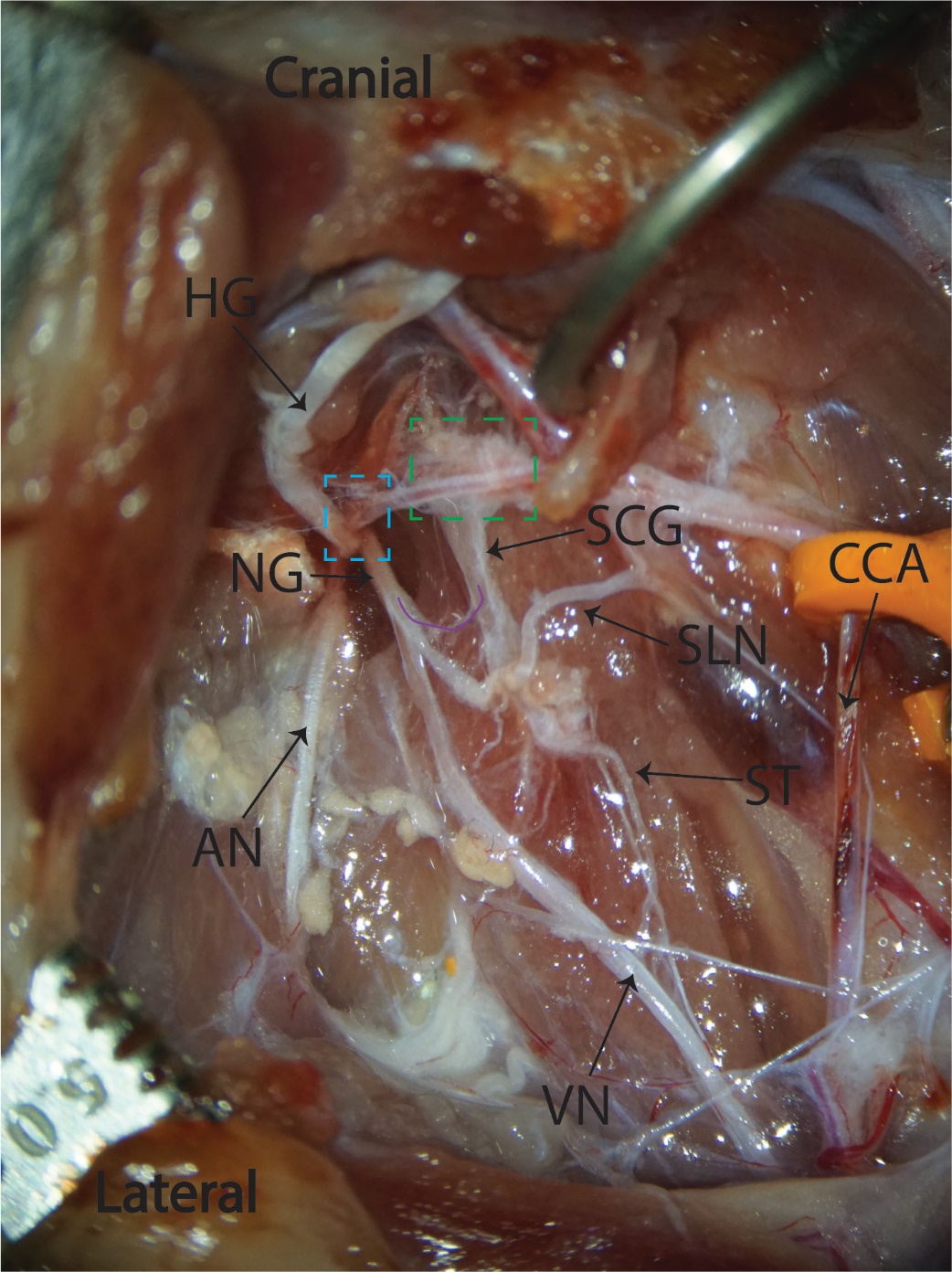

Supplementary Figure 4: **Cranial nerves in the cervical region near carotid bifurcation (right side)**. Cadaver dissections to study cranial nerves anatomy and their proximity to sympathetic system components. VN (VN, CN X), accessory nerve (AN, CN XI), Hypoglossal nerve (HG, CN XII), Superior laryngeal nerve (SLN), Superior cervical ganglia (SCG), Nodose ganglia (NG), common carotid artery (CCA), ST (ST).CN X, XI, XII all dived dorsal to enter the foramen (blue dotted square). There was a connecting branch between SCG and NG (purple solid line). Carotid bifurcation with connective tissue and plexus which was kept intact to preserve the location of carotid chemoreceptors, baroreceptors and other sensors reported at the carotid notch. There were branches from SCG joining the carotid plexus in the notch (green dotted square).

3) *Additional examples of the sympathetic trunk “hitchhiking” with the vagus nerve from different subjects. Note: Examples from both left and right side.*

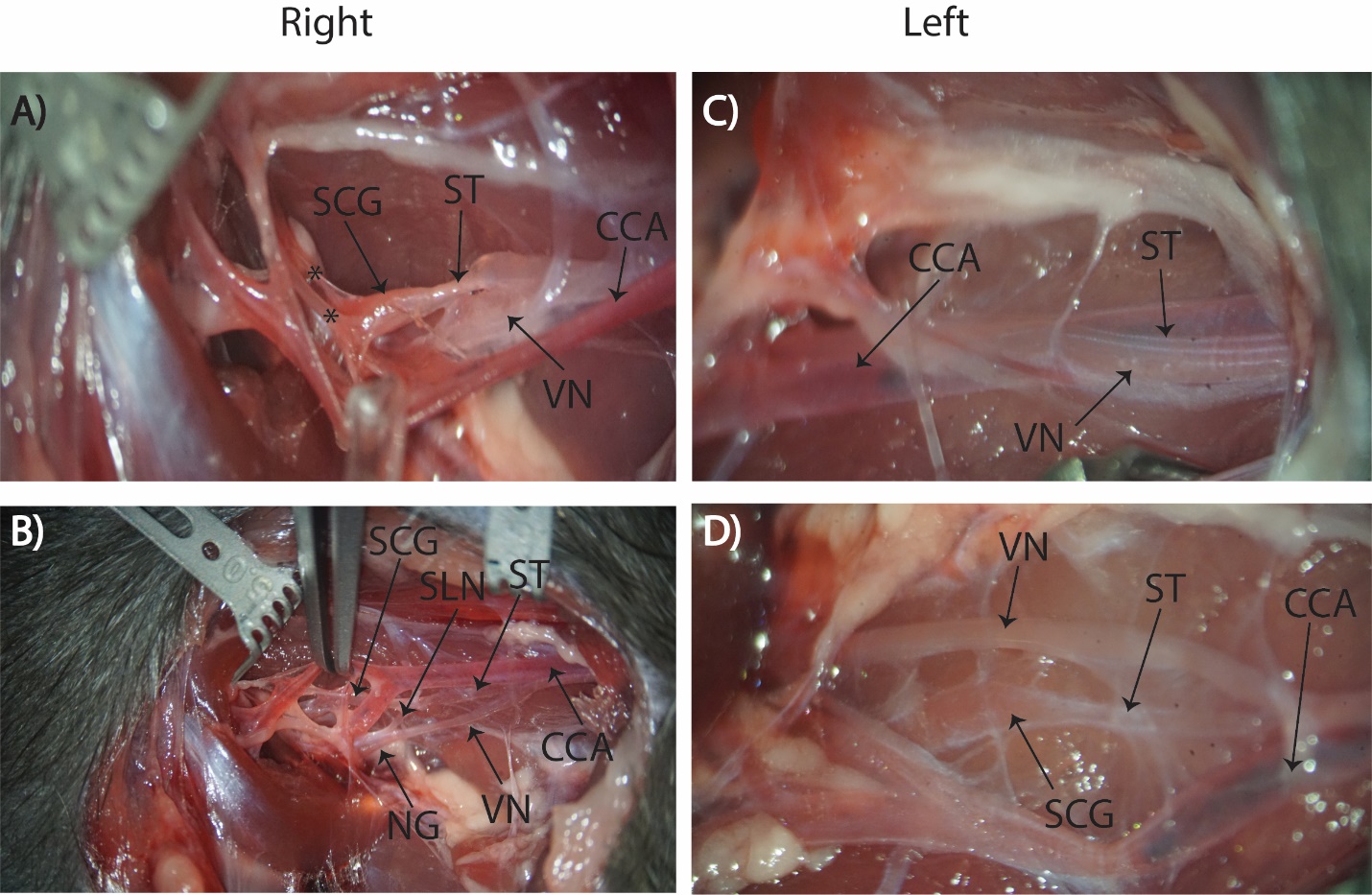

Supplementary Figure 5: **ST joint to VN running in parallel in carotid sheath.** A) ST originating from caudal pole of SCG and joining VN running together with common carotid artery. Multiple SCG branches joining ICA and carotid bifurcation. B) ST originating from caudal pole of SCG and VN originating from nodose ganglion (NG) with Superior laryngeal nerve (SLN) joining NG from medial to lateral side. ST and VN joining as they run caudal towards the sternum at the common cuff implantation location. C) VN and ST joint at the common cuff implantation location in an undissected carotid sheath. D) ST joining VN as traced caudal. Cross connecting branches between the two nerves along their cranial to caudal course.

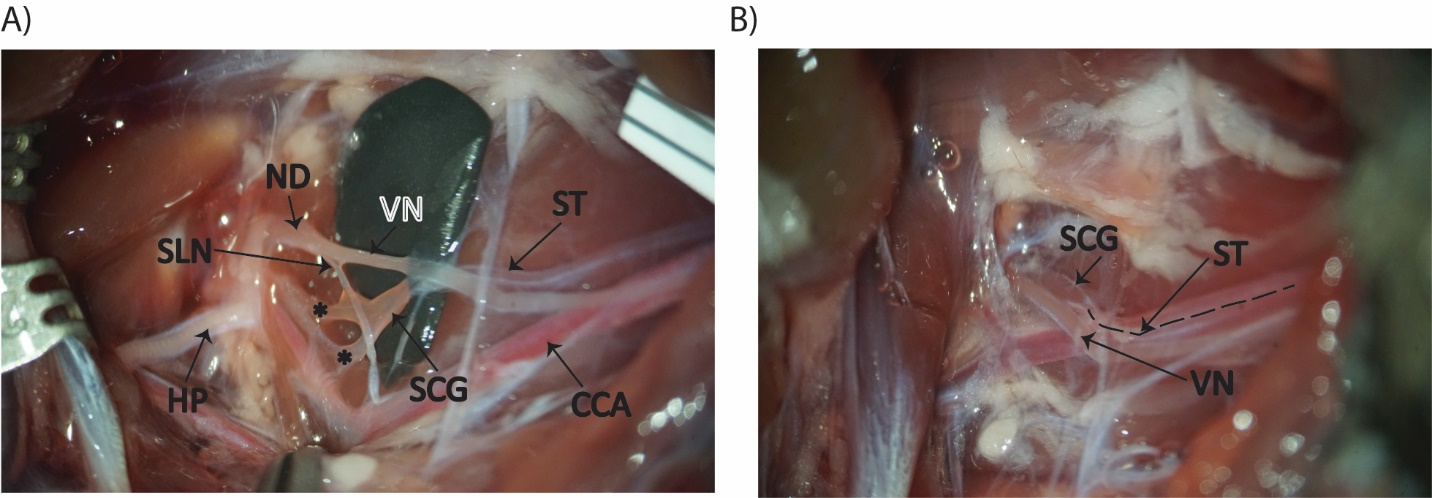

Supplementary Figure 6: **ST and VN running joint at one location but not throughout their course in the carotid sheath (Left side).** A) ST originating from SCG and VCN joining nodose ganglia (ND) with a crossover point between the two nerves as they run cranial to caudal. Superior laryngeal nerve (SLN) joining ND. Hypoglossal nerve (HP). B) SCG and ST joint to VN at cranial end of their course. ST and VN not joint as they run caudal but in close proximity on the same side of the carotid artery.

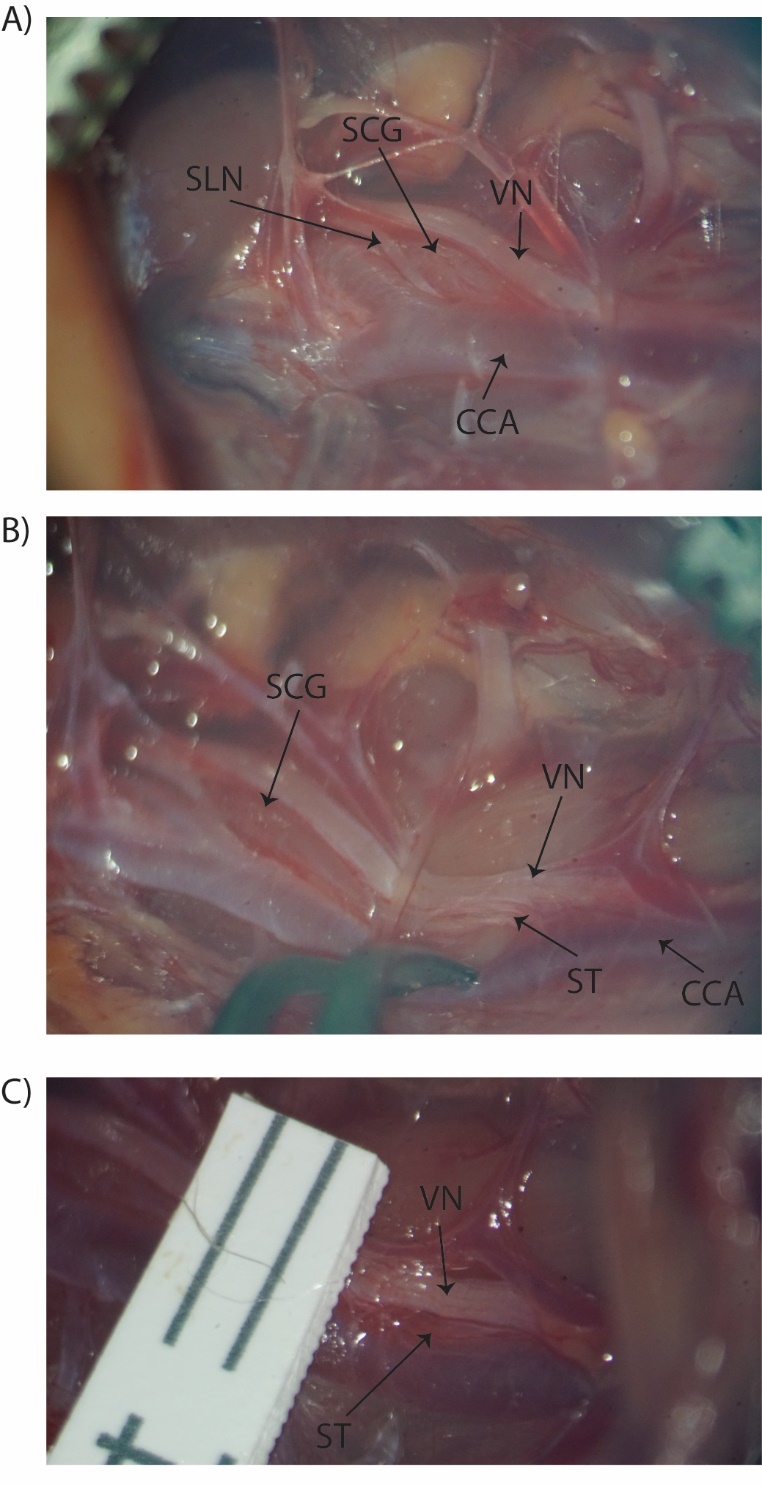

**Supplementary Figure 7: Left side ST and SCG joint to VN.** A) The SCG extending from dorsal side of the carotid bifurcation to the lateral side. It was found joint to the VN as VN dove dorsal into the foramen with SLN running over the SCG, under the bifurcation to join the VN at nodose ganglion. B) ST was found to be joint or ‘hitchhiking’ to VN and running on the lateral side of the common carotid artery. C) The ST was running dorsal to the VN still joint in the carotid sheath as the nerves were traced caudal towards the sternum from the bifurcation. Separating ST and VN revealed cross connections between the two nerves which had to be cut to isolate the two nerves for functional studies. The top two panels are from the same subject, and the bottom panel is from a different subject.

4) *Examples where the ST was not conjoint to the VN, but was in the carotid sheath on the opposite side of the VN.*

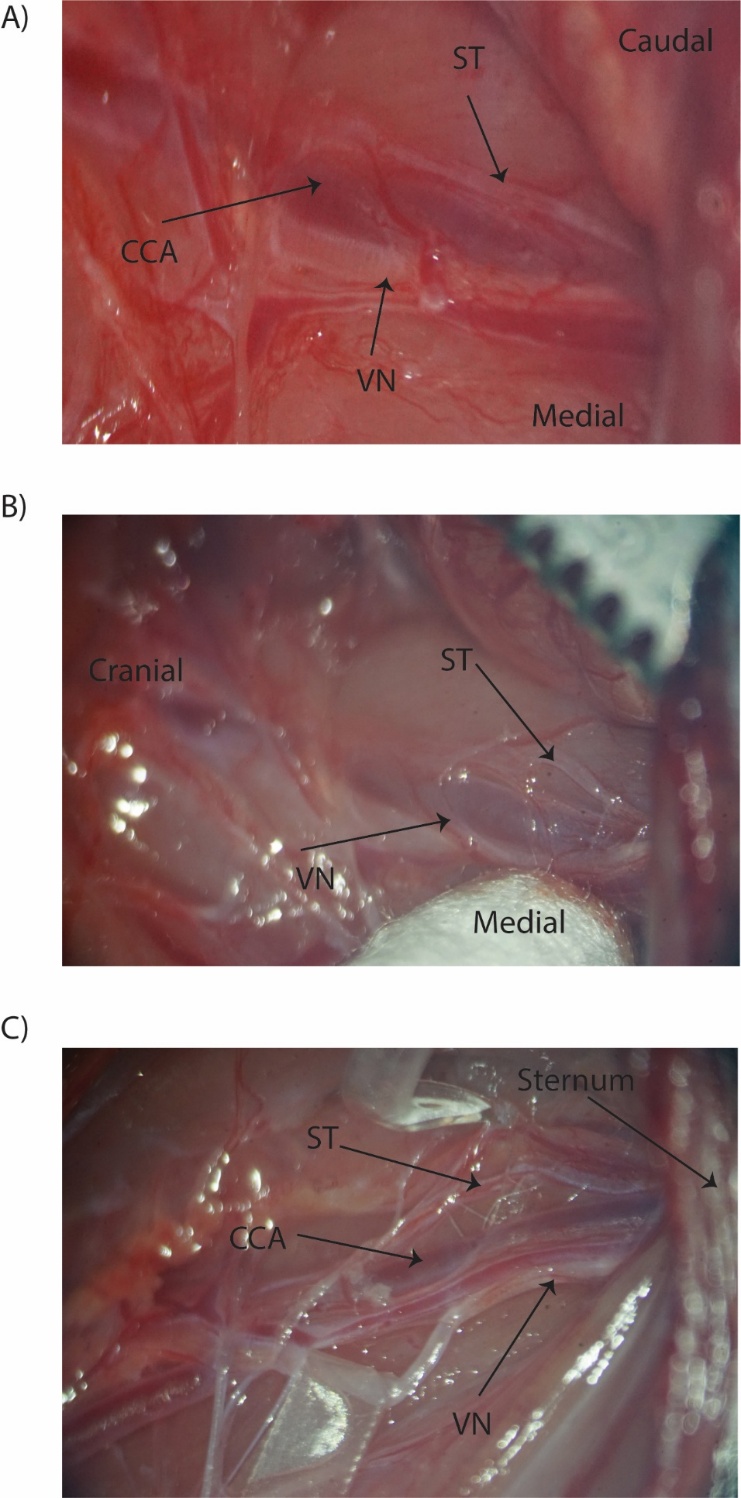

Supplementary Figure 8: **Left side ST was not joint to the VN (n=3).** A) In one animal, the ST and VN were found on the opposite sides, with VN medial and ST lateral of the carotid artery and not joint. They were easily isolated for electrode interfacing, however fine cross connections between the two nerves were found (not shown in figure). B) In a separate animal, ST was dorsal and lateral to the carotid artery at bifurcation and then coursed to the lateral side of the artery when traced caudal to the sternum. The VN was ventromedial to the carotid artery from the sternum, crossed over from medial to lateral of the artery before diving down. ST and VN were not joint and while their course crossed ~1mm caudal to the bifurcation, they were not joint. Cross connections were found between the two 1-2 mm caudal to the SCG.

5) *Additional subject example showing cross connecting branch between the ST and the VN along the Aortic depressor nerve along with the ST and the VN in the carotid bundle.*

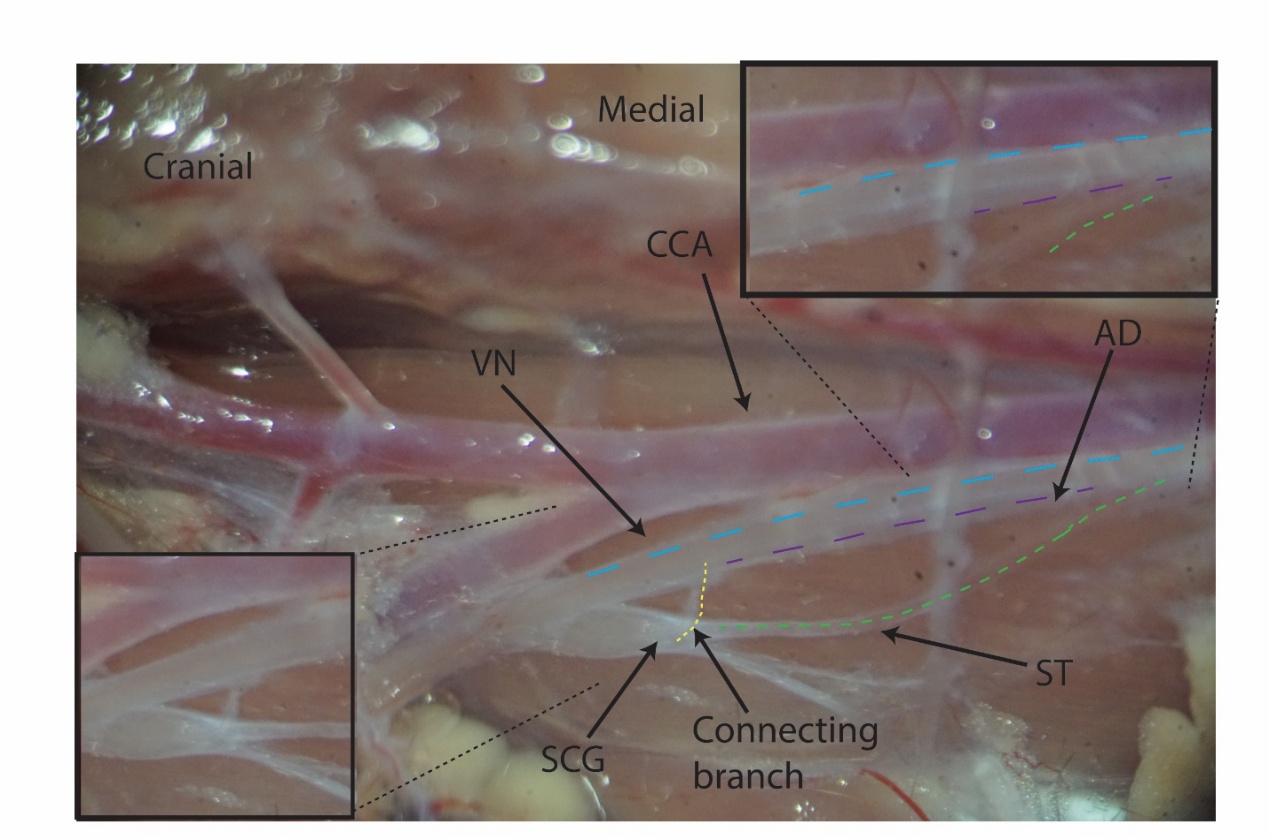

Supplementary Figure 9: **Cross connecting branch between VN and ST.** Cross connecting branch from caudal end of superior cervical ganglia (SCG) joining the VN when the ST and the VN are dissected out and separated. Aortic depressor nerve (AD) seen as independent nerve running parallel with the VN.

**Additional Functional data:**

6) *Cohort analysis for stimulation driven heart rate changes due to sympathetic trunk stimulation and vagus nerve stimulation from rodent left side (n=3).*

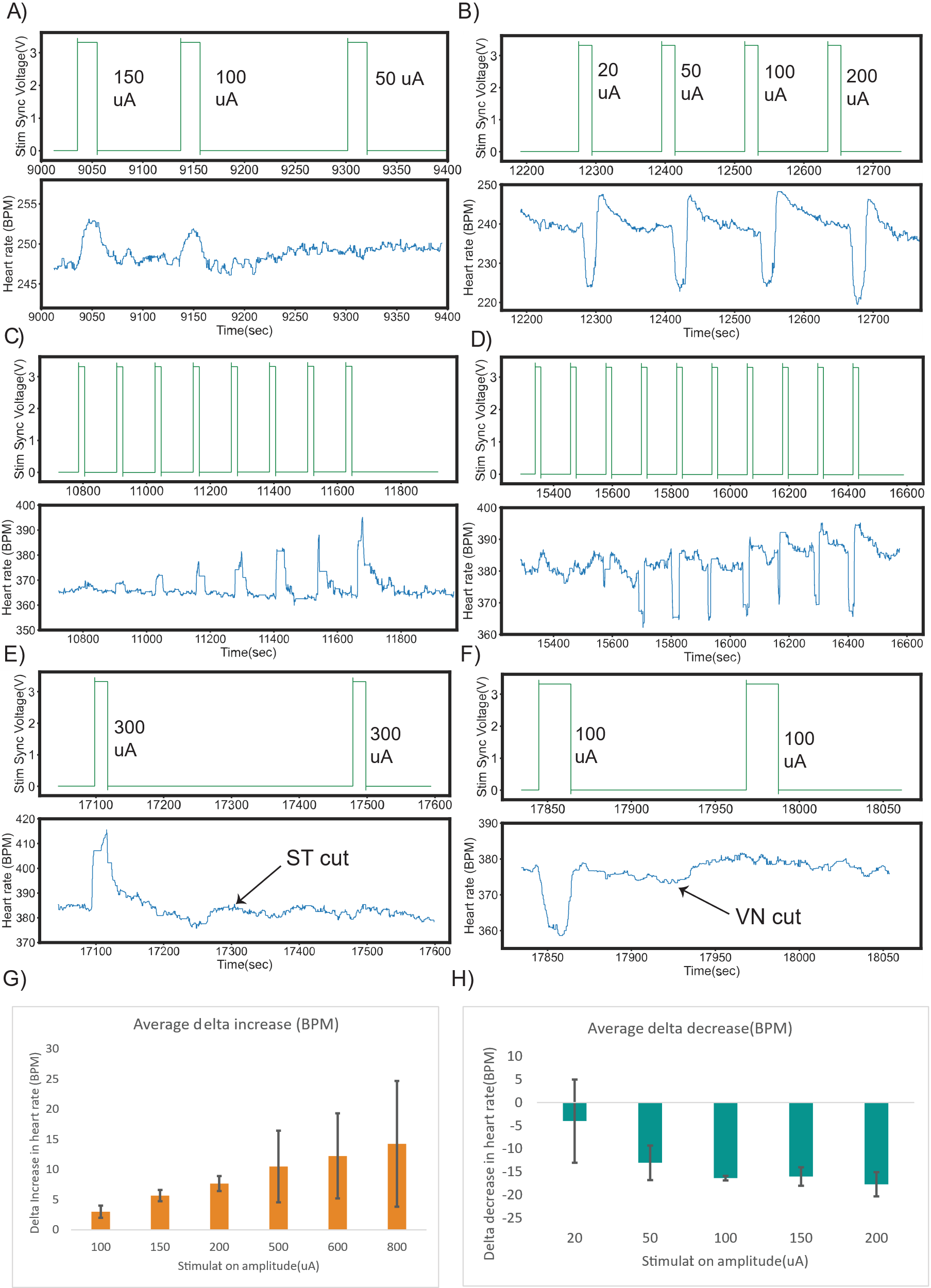

Supplementary Figure 10: **Dose response curves to study the effects of VN (VN) and ST (ST) stimulation on heart rate, left side(n=3).** A) Stimulation of isolated ST caused tachycardia B) Stimulation of isolated VN caused bradycardia C) Dose response curve for ST stimulation showing increased tachycardic response to increasing stimulation amplitude (6.25 Hz, 100-1000 uA) D) VNS with increasing stimulation amplitude showing slight tachycardia at lower stimulation amplitude (20 μA) with bradycardic responses at higher amplitudes (50-800uA) with a ceiling of 20 beats per minute (BPM). E) Cutting ST caudal to stimulation electrodes eliminated stimulation evoked tachycardia like right side verifying bilateral ST-tachycardia efferent circuits. F) Cutting VN caudal to stimulation electrodes got rid of stimulation evoked bradycardia G) Increasing ST stimulation amplitude (100-800 μ A) evoked tachycardia with maximum increase of 25 beats per minute (BPM) H) Increasing stimulation amplitude for VNS caused increasing bradycardic response till reaching max drop of ~20-25 at 200 μA which was 20 BPM less than that on right side which was 45 ± 32 BPM.

7) Temporal differences in HR to reach maximum delta change and return to baseline between VNS and STS

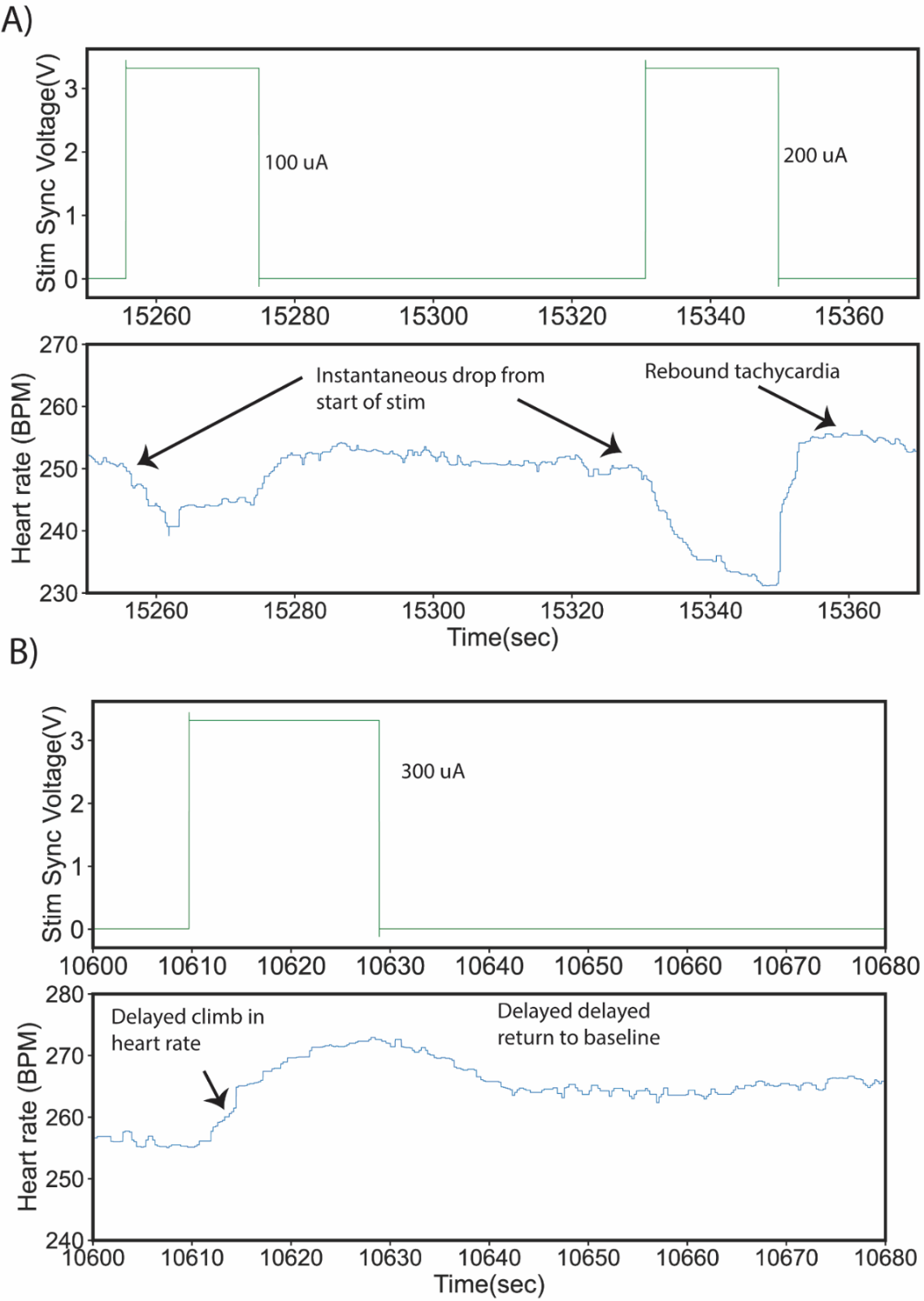

Supplementary Figure 11: **Representative animal responses showing the differences in temporal responses to stimulation for VNS and ST stimulation.** A) VNS caused an instantaneous drop with a rebound tachycardia and relatively quicker return to baseline B) ST stimulation caused a slow ramp increase of heart rate and a slow return to baseline pointing to possibly different mechanism of action for heart rate effects.

8) *Subject outlier in functional data omitted from cohort analysis*

**Spontaneous tachycardia in outlier subject:**

There were surgical complications in one animal during isolating and tracing the ST from SCG. A nerve branch from SCG (not the ST) was accidently transected. This resulted in spontaneous tachycardic responses [Supplementary figure 12]. These tachycardiac responses were observed before the ST was isolated and stimulated; so were not STS driven changes. The rest of the experimental protocol was followed as is, i.e. isolating and stimulating the ST and VN, independently and together. Stimulation of the isolated ST caused tachycardia and VNS caused bradycardia, as expected. However, STS driven tachycardia responses were in addition to the spontaneous tachycardia. Stimulating the VN and ST together resulted in a tachycardiac response at lower stimulation amplitudes, and stimulation at higher amplitudes resulted in a bradycardic response [Supplementary figure 12], which was the opposite of what was observed in the cohort. Transecting the ST caudal to the stimulation electrodes abolished stimulation-evoked tachycardia [Supplementary Figure 13], but not the spontaneous tachycardia. Transecting the VN eliminated stimulation evoked bradycardia as expected. The reasons for this outlier are difficult to disambiguate; whether they are due to stimulation evoked, the compromised physiological system due to surgical trauma, altered autonomic tone, or (most likely) a combination of all the above. The spontaneous tachycardia added another confound in analyzing this data, hence the animal was excluded from the cohort analysis.

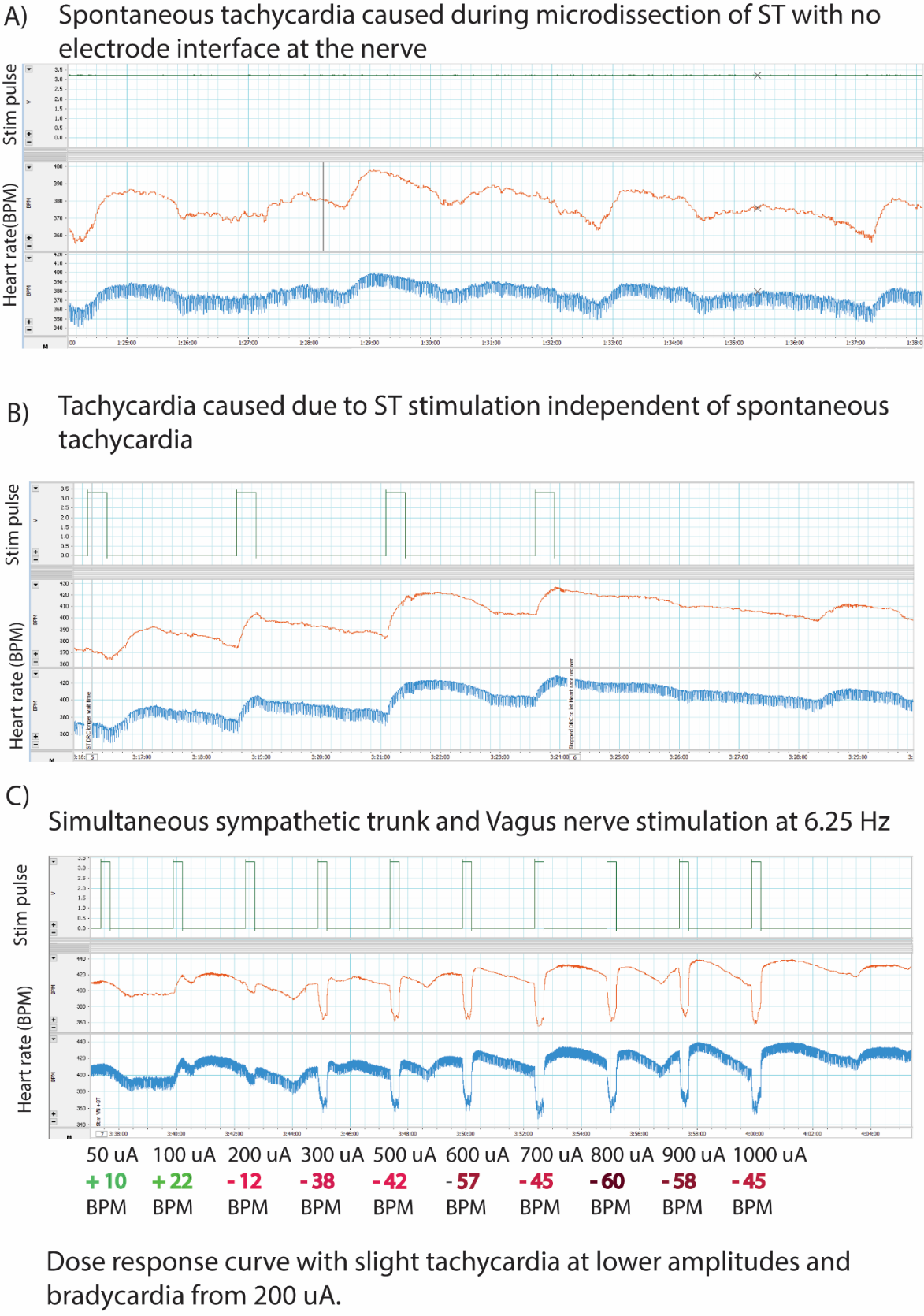

Supplementary Figure 12: **Spontaneous tachycardia independent of ST stimulation, outlier animal**. A) Spontaneous Tachycardia caused during microdissection with an increase of 20-30 beats per minute. B) ST stimulation evoked tachycardia was independent of spontaneous tachycardia. C) When stimulating both VN and ST simultaneous, tachycardia was caused at lower amplitudes and VN induced bradycardia at higher amplitudes.

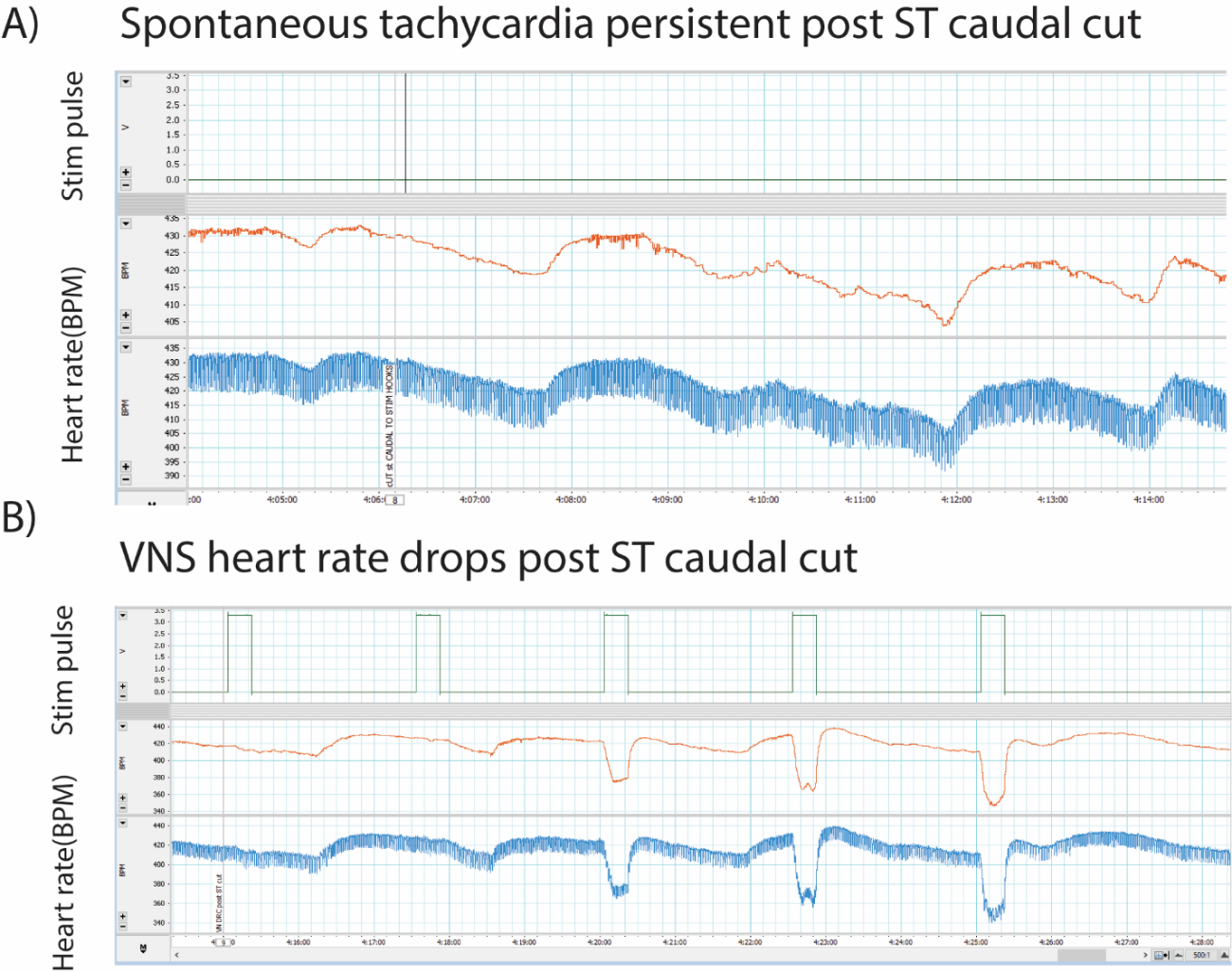

Supplementary Figure 13: **Spontaneous tachycardia outlier animal controls**. A) Spontaneous tachycardia persisted post ST cut caudal to stimulation electrode pointing to tachycardia caused independent of stimulation and through possible signaling pathway other than direct ST efferent control B) VNS caused bradycardia in the animal, however thresholds for bradycardia were slightly higher than the rest of cohort of subjects with bradycardia response starting at 200 μA. (Stimulation amplitudes: 50, 100, 300, 400, 500 μA)

**Controls:**

9) *Additional examples of ST and VN transections*

*Control 1: VN transection caudal to stimulation location eliminates bradycardic response*

*Control 2: ST transection caudal to stimulation location eliminates tachycardic response*

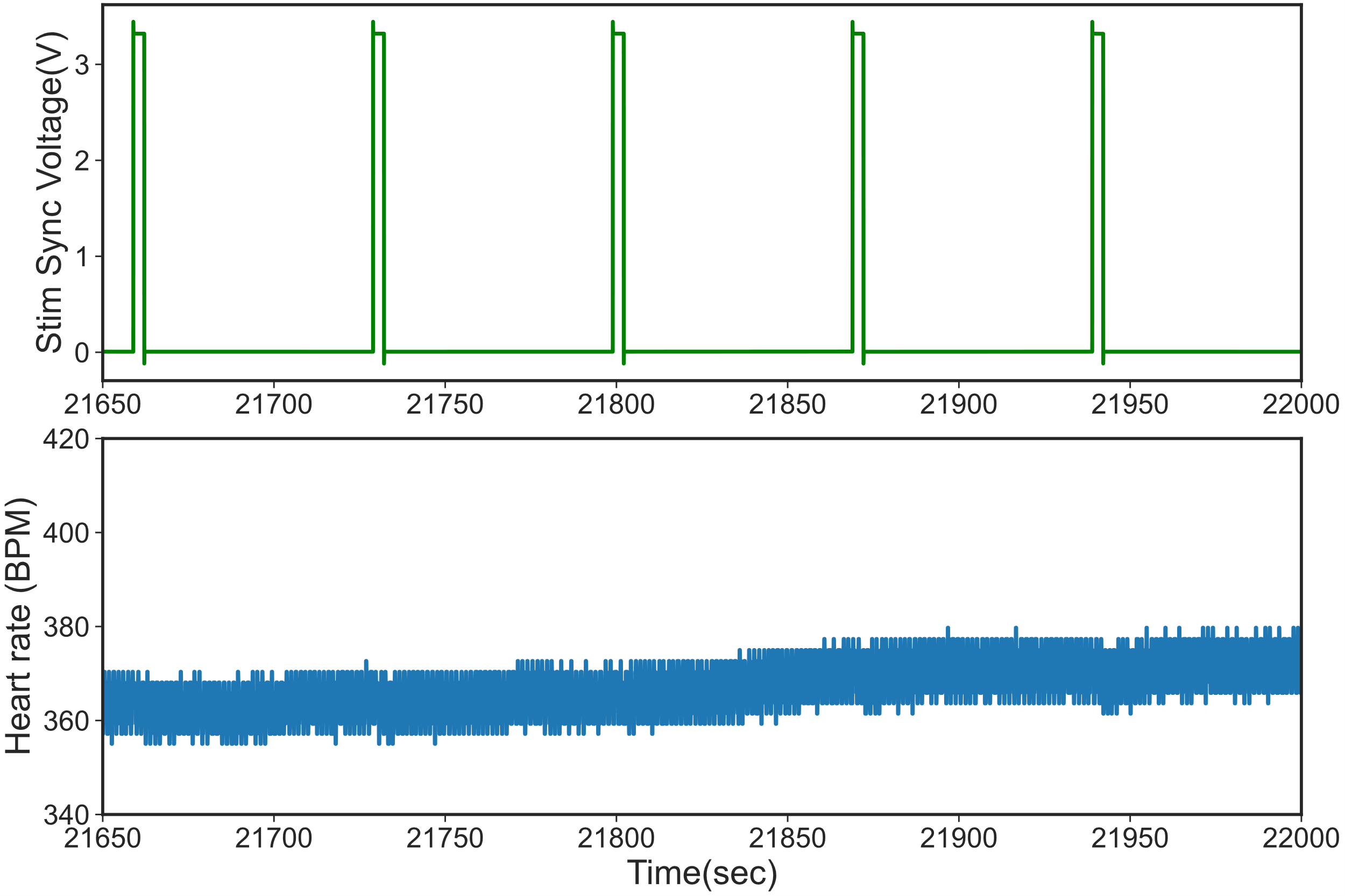

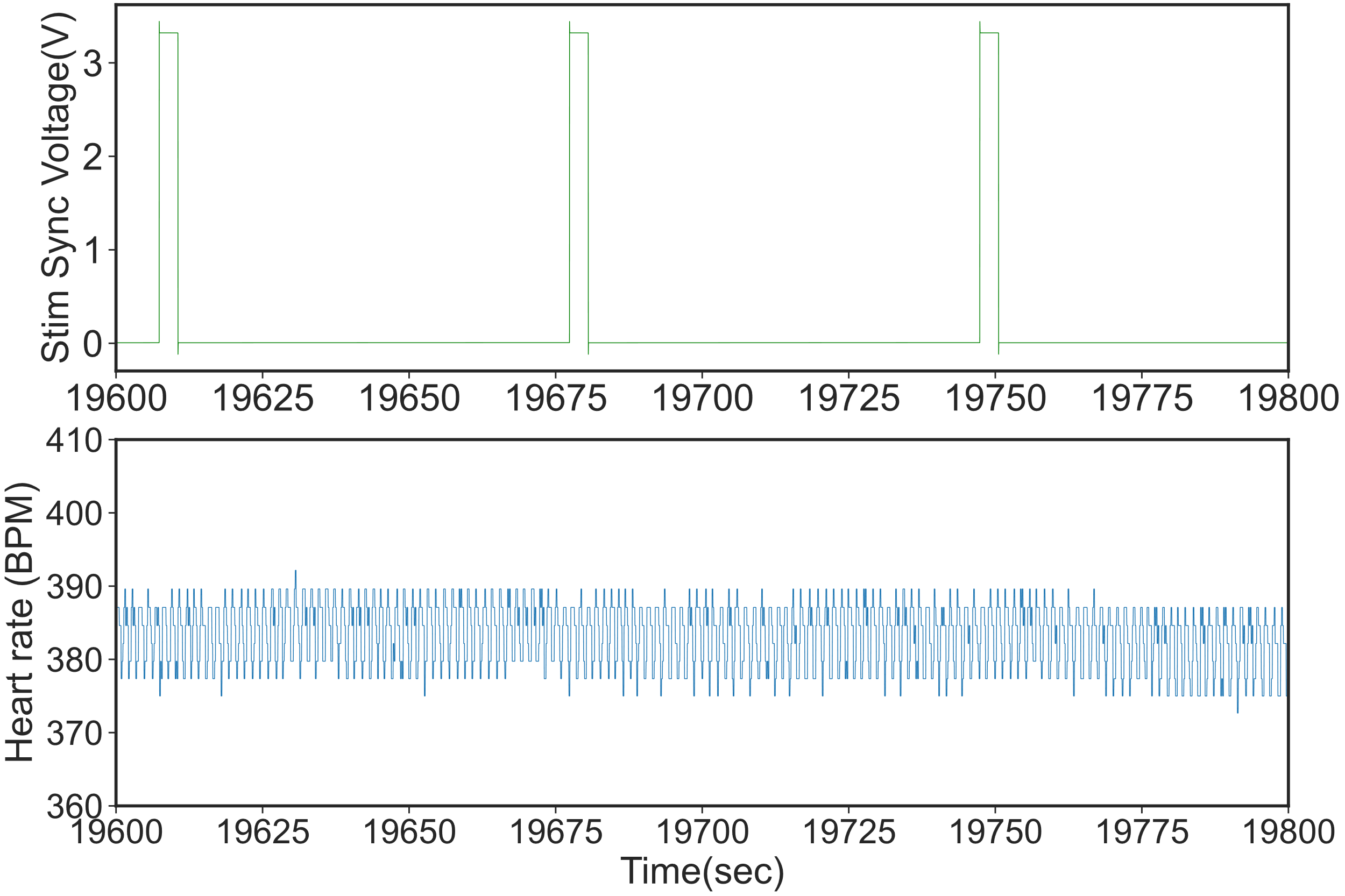

Supplementary figure 14: Loss of stimulation evoked heart rate changes post caudal cut to stimulation electrodes. Top) Loss of VNS evoked bradycardia post caudal cut. Bottom) Loss of ST stimulation tachycardia after cutting ST caudal to stimulation electrodes

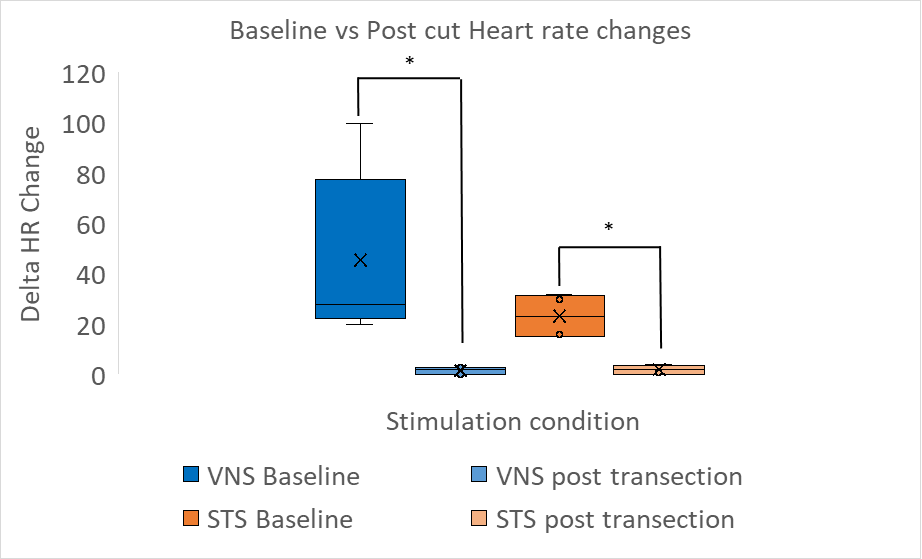

Supplementary figure 15: Box and Whiskers plot of delta HR changes pre and post nerve transections paired for stimulation amplitude (ex 200 uA stimulation pre and post transection) across multiple animals. VNS delta HR change baseline (Mean 46.6±33.30 BPM) compared to post transection (1.6±1.51 BPM), STS delta HR change baseline (Mean 23.25±8.99 BPM) compared to post transection (2±1.82 BPM).

10) *Stimulating the ST and the VN in conjunction yielded bradycardic response. Stimulating VN + STS yielded attenuated bradycardic responses as compared to VN alone in a subset of amplitudes tested in a dose response curve.*

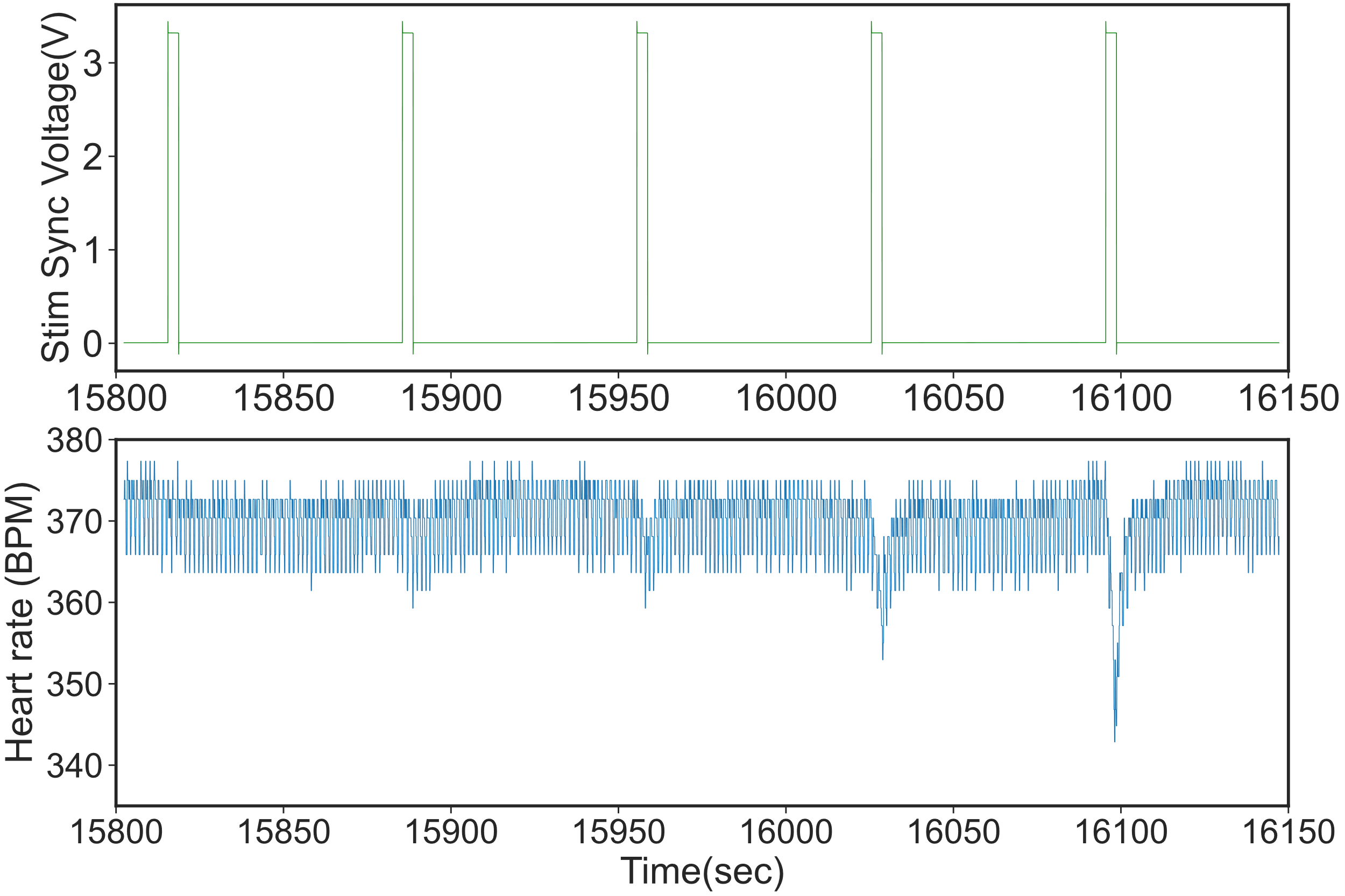

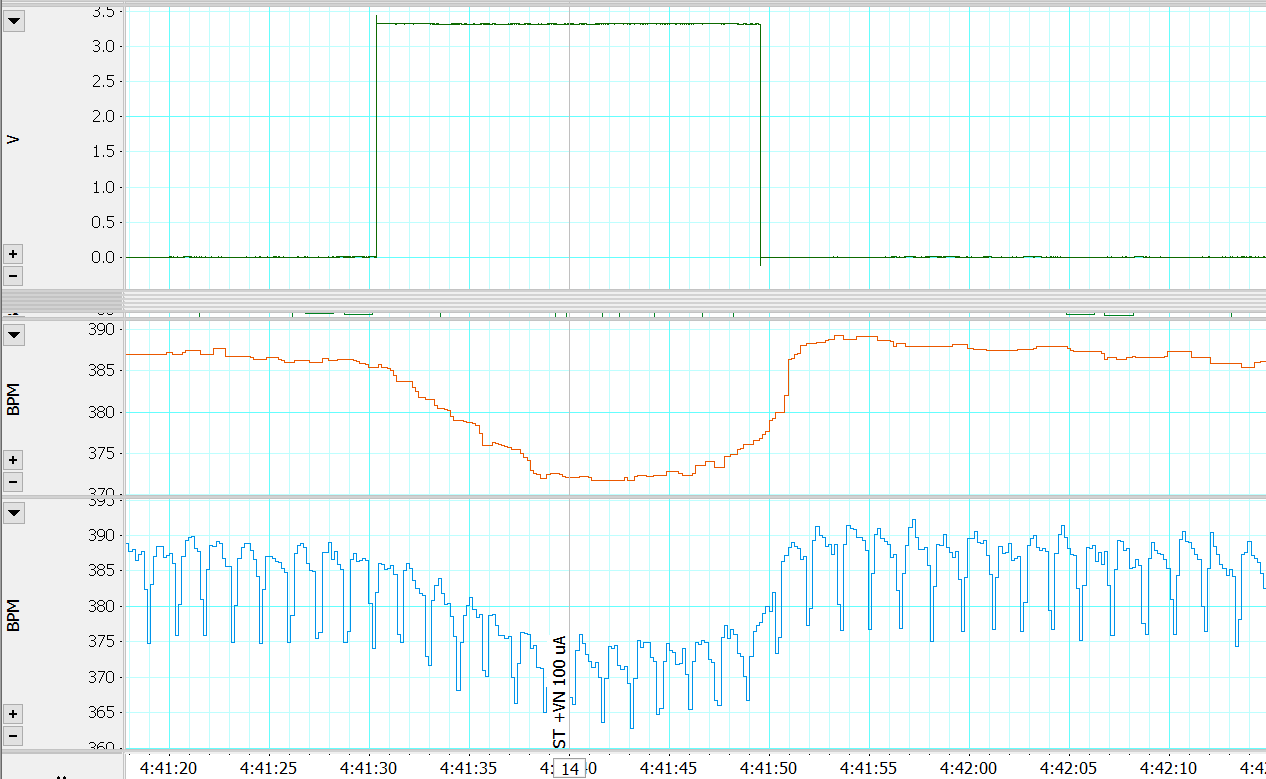

Supplementary Figure 16: **Dose response curve stimulation ST and VN together**. Top) Stimulation of ST and VN caused bradycardia with increasing stimulation amplitudes at 6.25 Hz, Stimulation amplitude (50,200,400,600,800 μA, right side) Bottom) Stimulation of ST and VN caused bradycardia (6.25 Hz,100 μA, left side)

Delta HR drop (BPM)

| Stimulation Amplitude (uA) | VNS | VNS+STS |
| --- | --- | --- |
| 100 | 25.7333 | 15 |
| 200 | 42.375 | 20 |
| 800 | 56 | 74.3333 |

Supplementary Figure 17: **Cohort comparison of HR drops for VNS alone and stimulating ST and VN together (n=4)**. Common stimulation amplitudes from dose response curves were selected such as there were a minimum of 3 subject responses for both groups with the same stimulation amplitude. The mean and standard deviation of the VNS +STS bradycardic response was found to be lower than VNS alone for 100 and 200 uA. However, it was not found to be statistically significant. Stimulation at 800uA resulted in increased bradycardic responses in STS+VNS as compared to VNS alone, which could be due to the reported phenomenon of anticipated antagonism. However, additional detailed studies are required to fully capture this phenomenon across stimulation amplitudes

11) *Clinically relevant frequencies 25 Hz and 40Hz tested for STS responses in a subset of animals.*

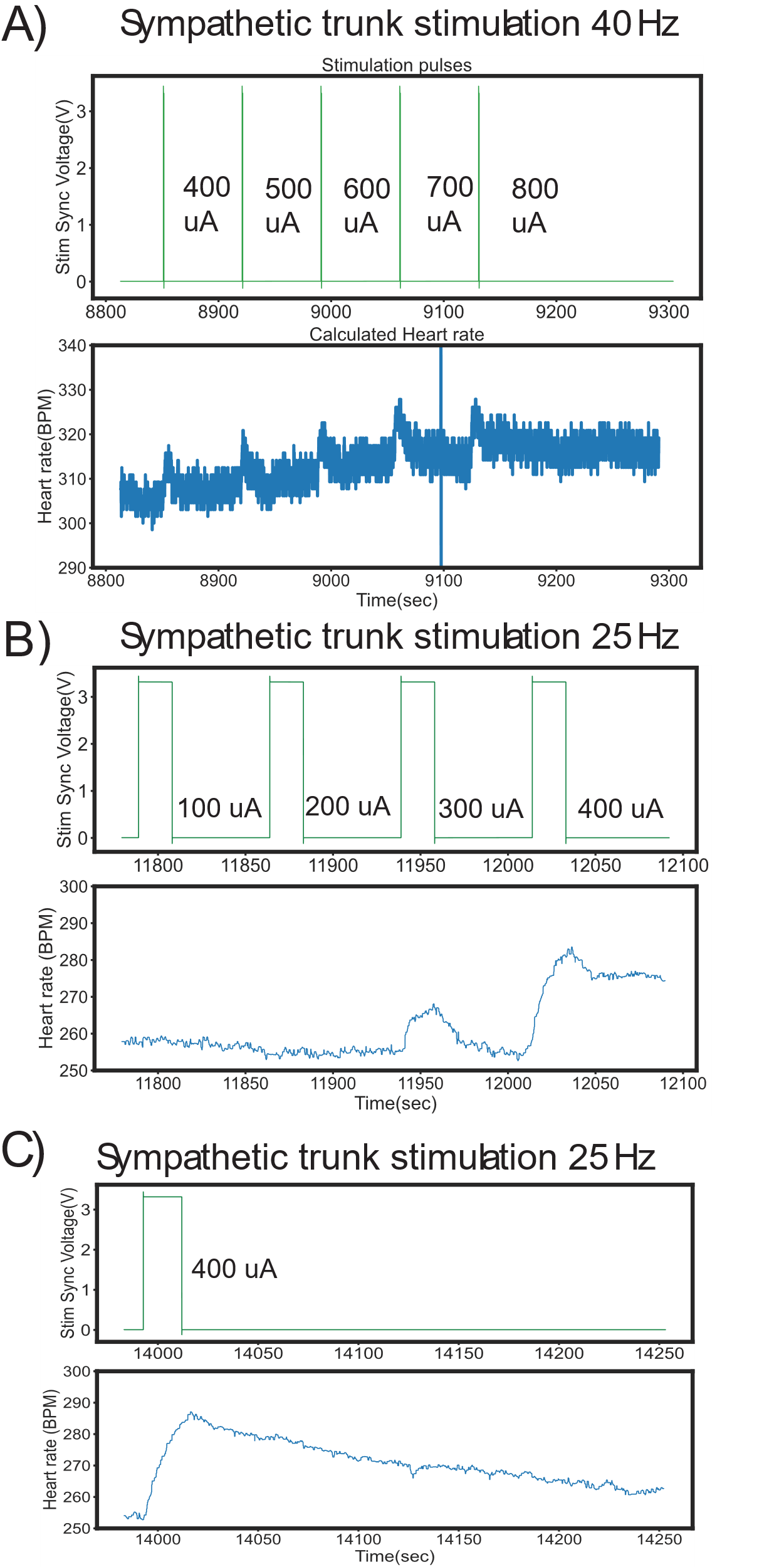

Supplementary Figure 18: **ST stimulation (STS) causes tachycardia independent of stimulation frequency.** Figures for showing STS caused tachycardia in subjects for clinically relevant stimulation frequencies of 25 and 40 Hz in a subset of animals.

**Histology:**

| **Figure 5 Panel** | **Distance between ST and VN** |
| --- | --- |
| A | 5 |
| B | 250 |
| C | 400 |
| D | 230 |
| E | 320 |
| F | 900 |

**Supplementary Table 1:** Tyrosine hydroxylase (TH+) post ganglionic sympathetic fibers bundle edge-to-edge distances to VN from histological samples in Figure 5.
